# Ribosome recycling factor ABCE1 depletion inhibits nonsense-mediated mRNA decay by promoting stop codon readthrough

**DOI:** 10.1101/870097

**Authors:** Giuditta Annibaldis, René Dreos, Michal Domanski, Sarah Carl, Oliver Mühlemann

## Abstract

Nonsense-mediated mRNA decay (NMD) is an essential post-transcriptional surveillance pathway in vertebrates that appears to be mechanistically linked with translation termination. To gain more insight into this connection, we interfered with translation termination by depleting human cells of the ribosome recycling factor ABCE1, which resulted in an upregulation of many but not all endogenous NMD-sensitive mRNAs. Notably, the suppression of NMD on these mRNAs occurs at a step prior to their SMG6-mediated endonucleolytic cleavage. Ribosome profiling revealed that ABCE1 depletion results in ribosome stalling at stop codons and increased ribosome occupancy in 3’ UTRs, indicative of enhanced stop codon readthrough or re-initiation. Using reporter genes, we further demonstrate that the absence of ABCE1 indeed increases the rate of readthrough, which would explain the observed NMD inhibition, since enhanced readthrough has been previously shown to render NMD-sensitive transcripts resistant to NMD by displacing NMD triggering factors like UPF1 and exon junction complexes (EJCs) from the 3’ UTR. Collectively, our results show that improper ribosome disassembly interferes with proper NMD activation.

**Highlights:**

- ABCE1 knockdown suppresses NMD of many NMD-sensitive mRNAs
- The observed NMD inhibition occurs at a stage prior to SMG6-mediated cleavage of the mRNA
- ABCE1 depletion enhances ribosome occupancy at stop codons and in the 3’ UTR
- ABCE1 depletion enhances readthrough of the stop codon
- Enhanced readthrough inhibits NMD, presumably by clearing the 3’ UTR of NMD factors

## INTRODUCTION

Translation, the synthesis of proteins by ribosomes according to the genetic information stored in messenger RNA (mRNA), is a central process in all forms of life. Although absolutely essential and hence requiring a high level of precision, translation involves many different factors that need to interact and function in a complex and highly orchestrated manner, and mistakes can occur at essentially every step. To achieve the necessary precision, cells have evolved numerous mechanisms to assess the integrity of mRNA and aberrancies in the functioning of ribosomes during translation (Lykke-Andersen and Bennett, 2014; Shoemaker and Green, 2012). In eukaryotes, nonsense-mediated mRNA decay (NMD) is one of these translation-dependent quality control pathways that degrades mRNAs that fail to properly terminate translation (Karousis and Muhlemann, 2019). In the past, NMD was believed to target and degrade exclusively mRNAs carrying an aberrant premature termination codon (PTC) that interrupts the open reading frame (ORF), thereby suppressing the production of potentially deleterious C-terminally truncated proteins (Behm-Ansmant and Izaurralde, 2006). However, with the advent of transcriptome-wide profiling methods it became clear that NMD regulates the levels of 5-10% of all mRNAs, the vast majority of them encoding perfectly functional full-length proteins (Colombo et al., 2017; Hurt et al., 2013; Mendell et al., 2004; Tani et al., 2012; Viegas et al., 2007; Wittmann et al., 2006; Yepiskoposyan et al., 2011). In the majority of these endogenous NMD targeted transcripts, there is a termination codon (TC) located >50 nucleotides upstream of the 3’-most exon-exon junction, a constellation that has been known for a long time to trigger NMD (Nagy and Maquat, 1998). Besides TCs located >50 nucleotides upstream of the 3’-most exon-exon junction, which can result from alternative pre-mRNA splicing, the presence of upstream open reading frames (uORFs), selenocysteine codons read as termination codons under low selenocysteine conditions, or the presence of introns in the 3’ untranslated region (3’UTR), long 3’ UTRs have also been documented to trigger NMD (Buhler et al., 2006; Eberle et al., 2008; Singh et al., 2008; Yepiskoposyan et al., 2011). Generally, the mRNA levels of NMD targeted transcripts with exon-exon junctions in the 3’ UTR are reduced by NMD to a greater extent than those with a long 3’ UTR, suggesting that exon junction complexes (EJCs) residing in the 3’ UTR of an NMD sensitive mRNA act as enhancers of NMD (Stalder and Muhlemann, 2008). EJCs are multiprotein complexes deposited 20-24 nucleotides upstream of exon-exon junctions during pre-mRNA splicing (Le Hir et al., 2001; Le Hir et al., 2000) and removed from the coding sequence during translation (Gehring et al., 2009), leaving behind only EJCs in the 3’ UTR on mRNAs undergoing translation. Notably, the transcriptome-wide studies also revealed that many pol II transcripts annotated as long noncoding RNAs are targeted by NMD (Colombo et al., 2017), in accordance with ribosome profiling studies showing that they in fact engage with ribosomes that translate short ORFs present in these transcripts (Calviello et al., 2016; Carlevaro-Fita et al., 2016; Ingolia et al., 2011).

The exact mechanism determining which RNAs will be targeted by NMD and which ones will be spared is subject to intense research and not yet well understood. The currently available data indicates that NMD ensues when a ribosome stalls at a termination codon (TC) for prolonged time because of a failure to properly or fast enough terminate translation (Amrani et al., 2004; Peixeiro et al., 2011). On mRNA with problems in translation termination, the so-called SURF complex (composed of SMG1, UPF1 and the release factors eRF1 and eRF3) was proposed to assemble at the TC (Ivanov et al., 2008; Kashima et al., 2006; Singh et al., 2008). SMG1-mediated phosphorylation of UPF1, a core factor of NMD, appears to be a key event in activating NMD, because hyper-phosphorylated UPF1 subsequently serves as a binding platform for the heterodimer SMG5/SMG7 and for the endonuclease SMG6, allowing for two different decay routes (Muhlemann and Lykke-Andersen, 2010). A third decay pathway operating via SMG5 and PNRC2 has also been proposed (Cho et al., 2013) but remains controversial (Loh et al., 2013; Nicholson et al., 2018). Moreover, a recent study challenged the SURF complex-based NMD activation model by showing that UPF3B rather than UPF1 interacts with the release factors (Neu-Yilik et al., 2017).

While EJC-enhanced NMD appears to be vertebrate-specific, extended 3’-UTR length has been reported as an NMD-inducing feature in a wide range of eukaryotes (Buhler et al., 2006; Eberle et al., 2008; Muhlrad and Parker, 1999; Pulak and Anderson, 1993; Singh et al., 2008). Consistent with the idea that ribosomes may fail to properly terminate translation when located too far away from the poly(A) tail, tethering of poly(A)-binding protein (PABP) nearby NMD-inducing TCs suppressed NMD in *S. cerevisae, D. melanogaster* and mammalian cells (Amrani et al., 2004; Behm-Ansmant et al., 2007; Eberle et al., 2008; Ivanov et al., 2008; Silva et al., 2008; Singh et al., 2008).

Prolonged ribosome stalling at NMD-inducing TCs increases the probability of readthrough, triggered by the accommodation of a near-cognate tRNA into the ribosome A-site, which then in turn can inhibit NMD (Hogg and Goff, 2010; Keeling et al., 2004) (Roy et al., 2015), presumably by stripping off the 3’ UTR NMD activators (e.g. UPF1, EJC) (Hogg and Goff, 2010). Frequent readthrough was shown to displace 3’ UTR-associated UPF1 from the mRNA, while inefficient readthrough allowed UPF1 (re-)association but blocked the initiation of mRNA decay at a subsequent rate-limiting step (Hogg and Goff, 2010). Although translation termination is generally kinetically favored over readthrough, the interaction between the TC and eRF1 is dependent on the nucleotide context, the eRF1 concentration and the availability of other factors needed for translation termination (Cridge et al., 2018). Besides the GTPase eRF3, the ribosome recycling factor ABCE1, an ATPase belonging to the ATP-binding cassette (ABC) protein family, plays crucial role in translation termination (Gerovac and Tampe, 2019). When an elongating ribosome arrives at a TC, eRF1 together with GTP-bound eRF3 binds the A-site of the ribosome. After eRF1-mediated hydrolysis and release of the nascent polypeptide chain from the tRNA in the P-site, ABCE1 interacts with eRF1 and promotes the splitting of the two ribosomal subunits and the release of the mRNA from the small subunit, providing the functional link between translation termination and a new round of translation initiation (Heuer et al., 2017; Mancera-Martinez et al., 2017). Several studies in different organisms demonstrated that silencing of ABCE1 affects the entire translation cycle and protein homeostasis of the cell (Guydosh and Green, 2014; Mills and Green, 2017; Sudmant et al., 2018; Young et al., 2015). In particular, the loss of ABCE1 led to increased ribosome stalling at TCs in an *in vitro* reconstituted translation system (Pisarev et al., 2010) and to the accumulation of post-terminating ribosomes in the 3’UTR of mRNAs in erythroid cells (Mills et al., 2016).

To gain more insight into the mechanistic links between translation termination and NMD, we investigated the effect of ABCE1 silencing on NMD. Here, we provide evidence that depletion of ABCE1 in HeLa cells not only leads to stalling of ribosomes at TCs but concomitantly also increases the rate of readthrough on these mRNAs. On many NMD-sensitive mRNAs, this increased readthrough suppresses NMD, consistent with the evidence that the ribosomes that translate into the 3’ UTR may prevent NMD by displacing NMD factors from these transcripts (Hogg and Goff, 2010). Thus, we provide evidence that ABCE1 and by inference proper ribosome recycling is required for NMD.

## RESULTS

### Knockdown of the ribosome recycling factor ABCE1 renders many endogenous NMD targets immune to NMD

Based on previous work indicating a link between translation termination and NMD (reviewed in (Karousis and Muhlemann, 2019), we hypothesized that interfering with translation termination would alter the abundance of many mRNAs, in particular NMD-sensitive transcripts. To test this hypothesis and identify the affected genes, we established an siRNA-based knockdown protocol that enabled us to reduce the protein level of the ribosome recycling factor ABCE1 to around 10% in HeLa cells (Fig. 1A). In three biological replicates, total RNA isolated from cells with an ABCE1 knockdown (ABCE1 KD) or cells with a control knockdown (CTRL KD) was depleted for rRNA, reverse transcribed and subjected to high throughput Illumina sequencing (RNA-seq) to assess the steady-state levels of mRNAs transcriptome-wide. Comparison of the three ABCE1 KD datasets among each other and with the three Ctrl KD datasets revealed an overall high correlation (Pearson correlation coefficients between 0.78 and 0.91), demonstrating not only good reproducibility among the biological replicates but also indicating that ABCE1 KD does not cause a global change in RNA levels (Fig. S1). Of the totally detected 47’830 transcripts, 3722 (7.8%) changed by 2-fold or more and with a *p*-value ≤ 0,05 (Fig. 1B). The detected changes occurred to a similar extent in both directions, with 1757 up- and 1965 down-regulated transcripts.

**Figure 1.**
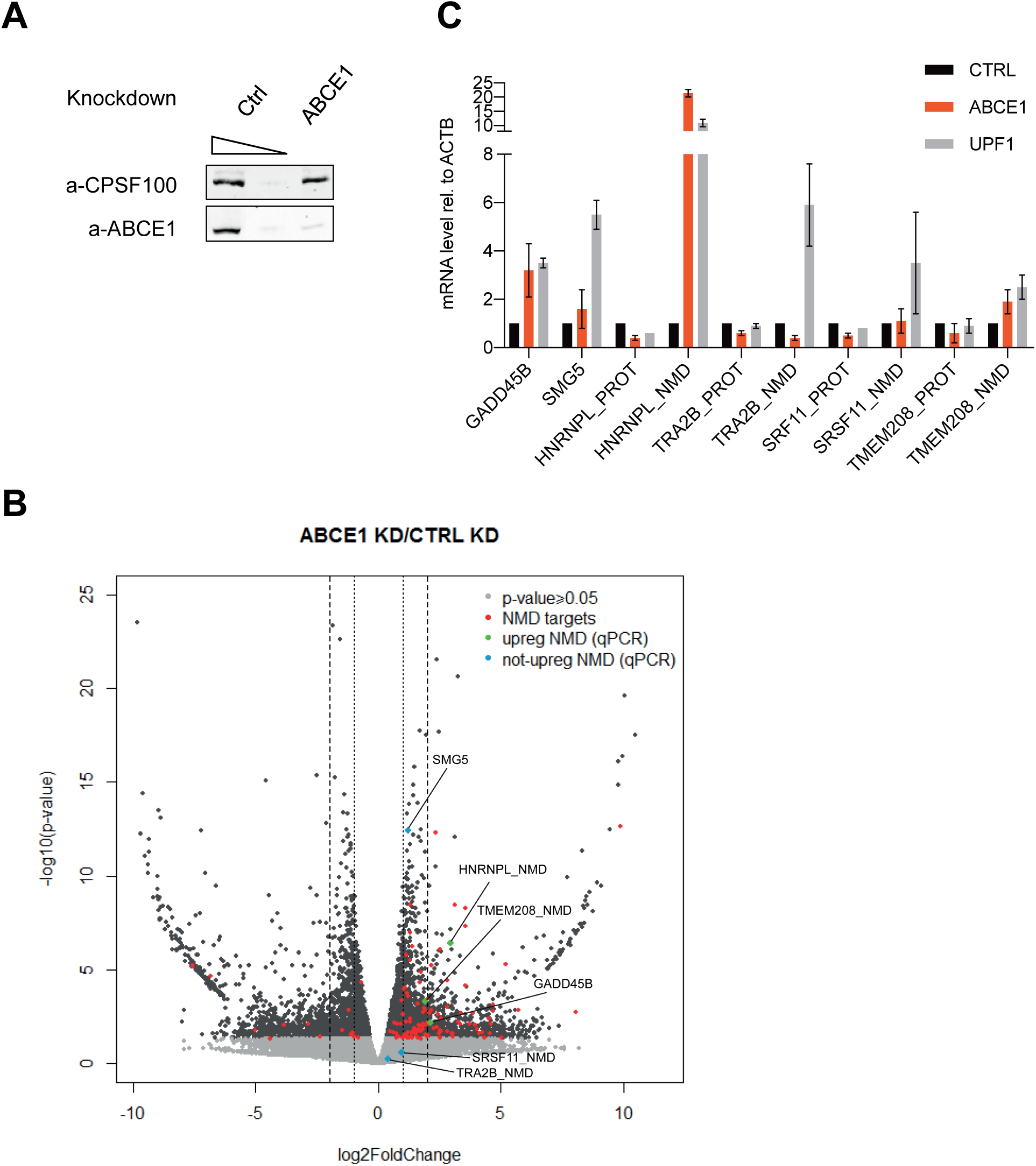
Depletion of ABCE1 inhibits NMD on many NMD sensitive transcripts. (A) Protein level analysis by western blot of ABCE1 knockdown in RNA sequencing samples. CPSF-100 serves as loading control. Of the Control KD, 100% and 10% of the sample was loaded, for better comparison of the ABCE1 signal with 100% of the ABCE1 KD sample. A representative blot of at least three independently performed experiments with similar results is shown. (B) Volcano plot showing changes in RNA levels from the RNA-seq data along the x-axis and *p*-values for each differential expression value along the y-axis. The average of three replicates for both conditions (ABCE1 KD and Control KD) is shown. The log2FoldChange of each transcript was calculated comparing ABCE1 KD against Control KD. Red dots correspond to high confidence NMD targets identified previously (Colombo et al., 2017), light grey dots show transcripts without significant changes (*p*-value ≥ 0,05). Labelled dots correspond to NMD targets validated by RT-qPCR in Figure 1C (blue, not upregulated transcripts; green, upregulated transcripts). The dotted and the dashed lines indicate log2FoldChanges of ±1 and ±2, respectively. (C) RT-qPCR measurements of well-characterized endogenous NMD sensitive transcripts under Control KD, ABCE1 KD or UPF1 KD. Mean values and standard deviation of mRNAs levels normalized to β-actin (ACTB) mRNA level are shown (n=4). NMD sensitive. splice isoforms are depicted as “NMD”, the corresponding protein coding isoforms as “PROT”

To assess the effect of ABCE1 KD on NMD-sensitive transcripts, we used a previously identified high-confidence set of endogenous NMD targeted genes (Colombo et al., 2017). This list comprises 1000 NMD targets that were identified based on their RNA level increase in UPF1, SMG6 and SMG7 knockdowns, as well as their decrease in the respective rescue experiments (Colombo et al., 2017). Of the 644 high-confidence NMD targets that we could detect in our analysis here, 109 (16.9%) changed significantly (*p*-value ≤ 0,05) by 2-fold or more upon ABCE1 KD, with the vast majority (96 transcripts) being upregulated (Fig. 1B, red and green dots). Thus, compared to non-NMD targets, the NMD-sensitive transcripts are overrepresented by a factor of 4 among the transcripts that increased in abundance upon ABCE1 KD, suggesting that ABCE1 KD inhibits NMD on a significant fraction of the NMD-sensitive transcripts.

To validate RNA-seq results, we used reverse transcription followed by quantitative PCR (RT-qPCR) to measure the relative RNA levels of a panel of well-characterized NMD-sensitive transcripts in total RNA prepared from HeLa cells with a knockdown of ABCE1, or a knockdown of the central NMD factor UPF1 as a positive control. For HNRNPL, TRA2B, SRSF11 and TMEM208, the NMD-sensitive transcript (labelled _NMD; Fig. 1C) arises from an alternative splicing event (intron retention, inclusion of a poised exon or usage of an alternative 5’ or 3’ splice site) as part of an autoregulatory feedback loop and the productively spliced form of the respective pre-mRNA encodes for the full length protein and is not targeted by NMD (labelled _PROT) (Li et al., 2017). We included RNA measurements of the _PROT splice forms of these genes as a control and to exclude that the increases observed for the _NMD splice forms were due to changes at the level of transcription or splicing. The RT-qPCR results were in good agreement with the RNA-seq data, confirming the upregulation of a fraction of the NMD-sensitive transcripts (GADD45B, HNRNPL_NMD, and TMEM208_NMD) in ABCE1 KD, while another fraction of NMD-sensitive transcripts (TRA2B_NMD and SRSF11_NMD) was not affected and SMG5 was weakly but not statistically significantly upregulated. Altogether, these data indicated that ABCE1 KD inhibits NMD on a sub-population of NMD-sensitive transcripts, but none of the known NMD-inducing features could readily explain why certain NMD targets were up-regulated by ABCE1 KD while others remained unaffected.

### ABCE1 knockdown also inhibits NMD of well-studied NMD reporter constructs

Since only a sub-population of endogenous NMD targets were upregulated by ABCE1 KD, we wondered whether widely used NMD reporter constructs were also affected by ABCE1 depletion. Therefore, we performed the ABCE1 knockdown in HeLa cells that stably express either of two well-characterized NMD reporters, namely the TCRβ (Mohn et al., 2005) or immunoglobulin-μ (Ig-μ) minigenes (Buhler et al., 2004). Of both reporter genes, there exists a PTC-free (so called “wild-type”) version that produces an mRNA which is not sensitive to NMD (TCRβ WT and miniμ WT), and a version harbouring a PTC in the middle of the coding sequence (TCRβ ter68, miniμ ter310) that strongly triggers NMD (Fig. 2A, B). Again, or siRNA-based knockdown protocol resulted in a reduction of ABCE1 protein to about 10% of the normally present amount (Fig. 2A, B). Measurements of the respective relative reporter RNA levels by RT-qPCR showed that both NMD-sensitive transcripts increased by 2.5-fold (TCRβ ter68) and 8-fold (miniμ ter310) upon ABCE1 KD, whereas the PTC-free versions remained unchanged (miniμ WT) or even slightly decreased (TCRβ WT) (Fig. 2C, D), indicating that NMD of these two reporter transcripts was inhibited by ABCE1 KD. Hence, we reasoned that these reporters would be useful to further investigate how ABCE1 inhibited NMD.

**Figure 2.**
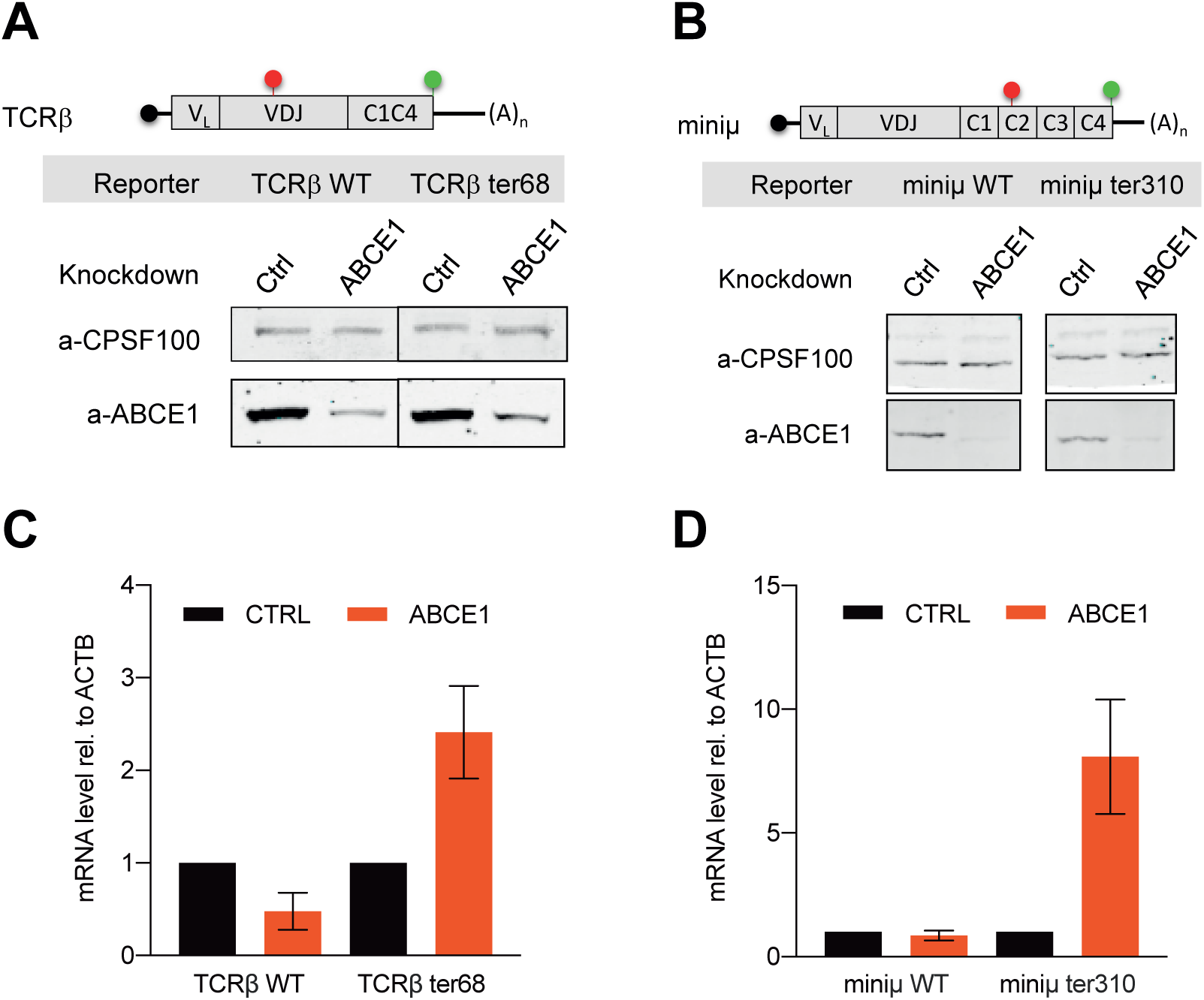
Depletion of ABCE1 increases mRNA levels of NMD sensitive reporters. (A, B) Schematic representation of TCRβ and miniµ reporter mRNAs. Red dots indicate the position of premature termination codons (ter68 in TCRβ and ter310 in miniμ), green dots represent the normal termination codon (top). Western blot analysis monitoring efficiencies of ABCE1 depletion in HeLa cells stably expressing TCRβ WT or ter68, and miniμ WT or ter310, respectively (bottom). CPSF-100 served as loading control. Representative blots of at least three independently performed experiments with similar results are shown. (C, D) RT-qPCR of HeLa cells expressing TCRβ or miniμ reporters. Mean values and standard deviations of mRNAs levels normalized to β-actin (ACTB) mRNA levels were calculated (n=4).

### ABCE1 knockdown inhibits NMD upstream of SMG6-mediated endonucleolytic cleavage on NMD-sensitive reporters

The observed NMD inhibition of the two NMD reporters TCRβ ter68 and miniμ ter310 prompted us to further investigate at which point in the NMD pathway this inhibition occurs. In yeast, depletion of the ABCE1 homolog Rli1 has been shown to lead to the accumulation of a 3’ RNA decay fragment of an NMD-sensitive mRNA as a consequence of a defect in translation termination (Serdar 2017). The authors of this study speculated that in the absence of Rli1, stalled ribosomes at the premature stop codon would block the 5’-3’ exonuclease Xrn1, resulting in the accumulation of this 3’ RNA decay intermediate. To see whether ABCE1 KD would also cause the appearance of a similar decay intermediate in human cells, we examined by northern blot WT and PTC-containing TCRβ and miniμ transcripts expressed in HeLa cells (Fig. 3). As documented by the western blots, a reduction to about 10% of the normal ABCE1 protein levels was achieved in the ABCE1 KD (Fig. 3 A, B). For both transcripts, we used probes that hybridize to the RNA downstream the PTC, which allows the detection of both the full-length transcript and the putative 3’ fragment (Fig. 3 C, D, top). While the NMD-insensitive WT versions of TCRβ and miniμ mRNAs did not change significantly as expected, we detected a marked increase of the full-length transcripts of the NMD-sensitive versions (TCRβ ter68 and miniμ ter310) upon ABCE KD (Fig. 3 C, D, bottom). However, no 3’ decay intermediates were detected, indicating that in human cells, ABCE1 depletion inhibits the NMD pathway upstream the SMG6-mediated endonucleolytic cleavage of the RNA.

**Figure 3.**
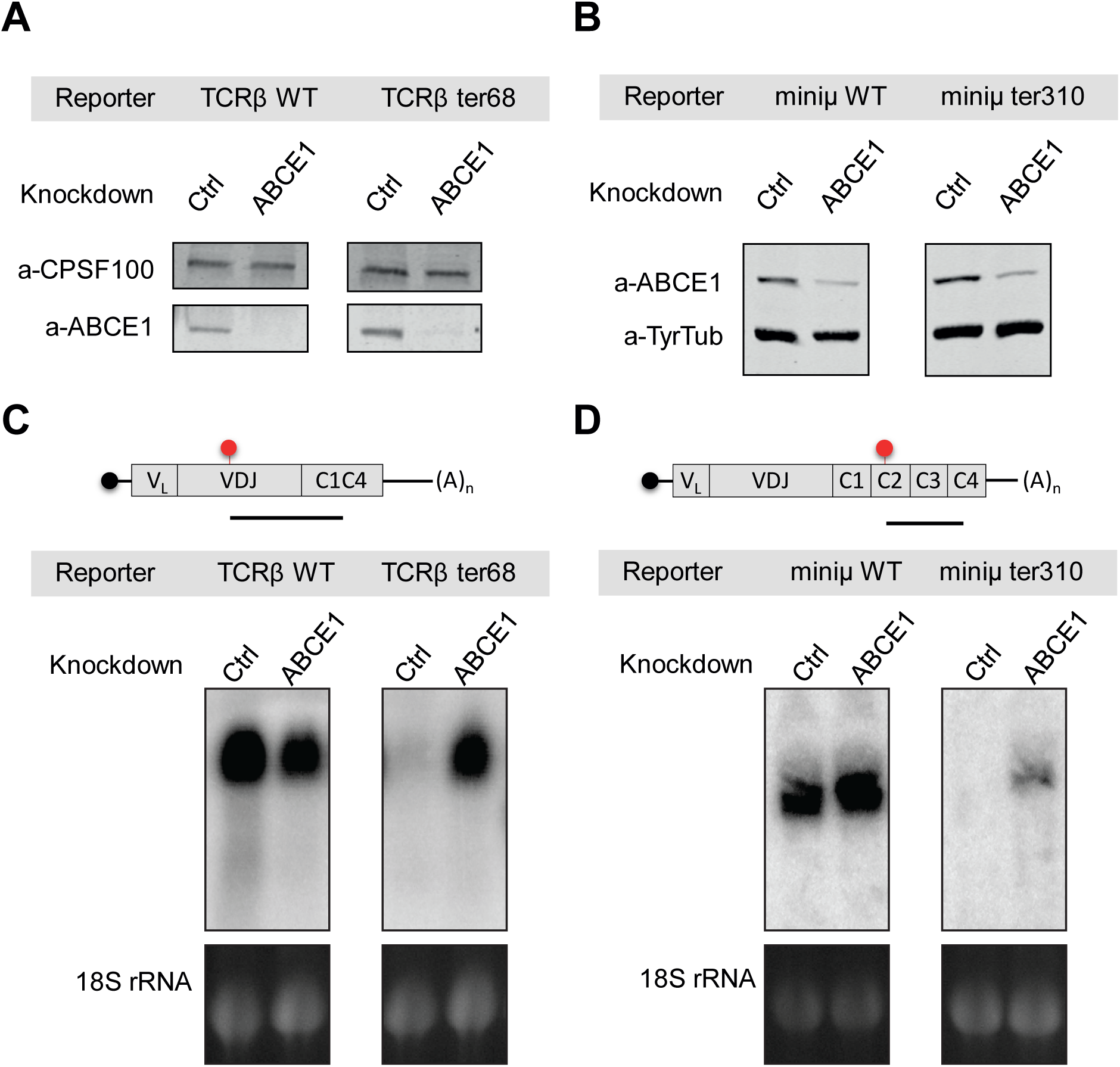
ABCE1 knockdown inhibits NMD upstream of SMG6-mediated endo-nucleolytic cleavage on NMD-sensitive reporters. (A, B) Western blot analysis monitoring efficiencies of ABCE1 depletion in HeLa cells stably expressing TCRβ or miniμ NMD reporters or NMD insensitive WT, repectively (top). CPSF-100 or Tyrosine Tubulin (TyrTub) served as loading control. The blots are representative of at least two independently performed experiments with similar results. (C, D) Schematic representation of TCRβ and miniμ reporters. Position of the probes used for northern analysis is indicated as a black line below the constructs (top). Northern blot analysis of total RNA isolated from HeLa-TCRβ (WT or ter68) or from HeLa- miniμ (WT or ter310) cells depleted for ABCE1 (bottom). 18S rRNA levels detected from the agarose gel stained with ethidium bromide serves as loading control.

### NMD sensitive mRNAs are strongly associated with polysomes in ABCE1 depleted cells

Since NMD is dependent on active translation (Carter et al., 1995; Thermann et al., 1998), we next examined whether ABCE1 depletion might inhibit NMD indirectly by causing a global translation inhibition. Notably, a previous study has reported that ABCE1 KD in human cells reduces the general translation level (Toompuu et al., 2016). To assess the overall translation activity in control and ABCE1 KD cells, we pulse-chased HeLa cells with puromycin and detected puromycylated nascent polypeptide chains using an antibody against puromycin (Fig. S2). The western blot results revealed a decrease of total protein synthesis by about 40% in cells depleted of ABCE1. However, this overall slight reduction in protein synthesis is unlikely to account for the selective inhibition of NMD on a fraction of the NMD targets, and moreover the puromycylation assay does not address which step of translation is affected by ABCE1 KD. There is evidence that ABCE1 depletion causes ribosome stalling at termination codon and an increased ribosome occupancy in the 3’UTR, suggesting mainly an impairment of translation termination (Mills et al., 2016; Sudmant et al., 2018; Young et al., 2015). According to this model, mRNAs are expected to still be associated with polysomes upon ABCE1 KD, and strong association of mRNAs with polysomes would provide indirect evidence that these mRNAs are still actively translated. To test this hypothesis, we knocked down UPF1 or ABCE1 in HeLa cells (Fig. 4A) and performed polysome profiling (Fig. 4B) to assess the association of different transcripts with monosomes (M) and polysomes (P) (Fig. 4C). The relative distribution between the monosome and the polysome fractions were quantified by RT-qPCR for two NMD-sensitive (HNRNPL_NMD, TMEM208_NMD) and two NMD-insensitive (HNRNPL_PROT, TMEM208_PROT) endogenous transcripts. Remarkably, compared to the control, the association of the NMD-sensitive transcripts HNRNPL_NMD and TMEM208_NMD with polysomes was increased under ABCE1 KD or UPF1KD conditions, similar to the protein coding isoforms that are actively translated under control and KDs conditions. We conclude that, even though ABCE1 KD slightly reduces overall the translation rate in cells, this cannot account for the inhibition of NMD on these transcripts, since they are still actively translated judged by their association with polysomes.

**Figure 4.**
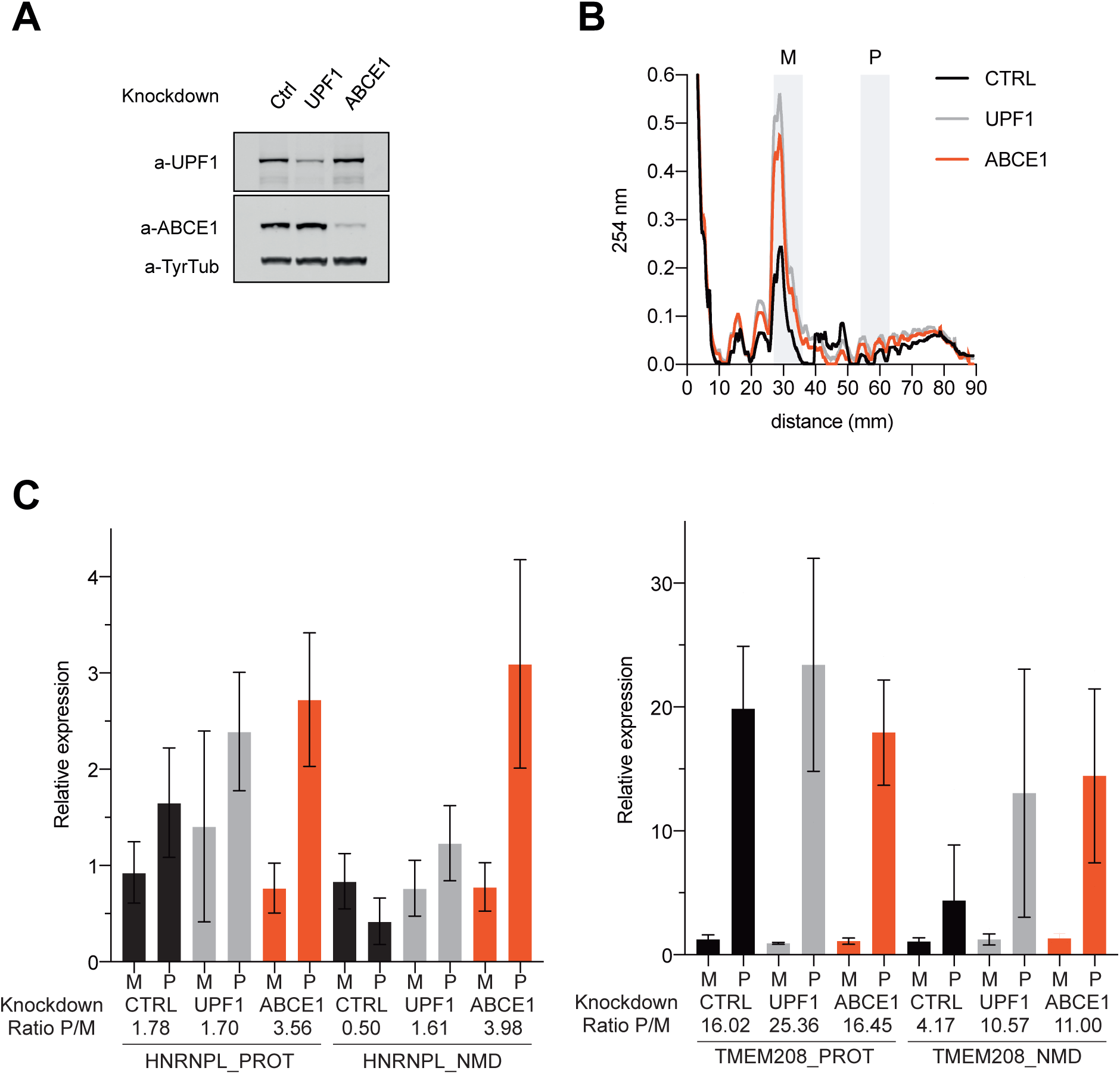
NMD sensitive mRNAs are strongly associated with polysomes in ABCE1 depleted cells. (A) Western blot of HeLa lysates to assess the ABCE1 and UPF1 knockdown efficiencies in the samples used for the polysome profiling experiments. Tyrosine Tubulin (TyrTub) served as loading control. The blot is representative of at least three independently performed experiments with similar results. (B) Representative polysome profiles of Control KD, ABCE1 KD and UPF1 KD samples. Similar results were obtained in three independently performed experiments. Fractions corresponding to monosomes (M) or polysomes (P) used for RNA isolation and RT-qPCR analysis are highlighted in light grey. (C) Quantification by RT-qPCR of NMD sensitive (NMD) or insensitive (PROT) isoforms of HNRNPL and TMEM208 RNAs are shown. Bars indicate the fraction of the mRNA in monosomes (M) and polysomes (P) under different knockdown conditions. Relative ratios between polysome and monosome fractions are shown. Mean values and standard deviations of mRNA levels relative to first monosome fraction were calculated with (n=3).

### Depletion of ABCE1 induces ribosome stalling at termination codon and an enrichment in 3’ UTR ribosome occupancy

To gain more insight about translation changes and ribosome occupancy at TCs in the absence of ABCE1, we performed ribosome profiling in HeLa cells in which ABCE1 was knocked down as described before (Fig. S5A) using a modified version of the protocol published by Ingolia and coworkers (Ingolia et al., 2012). In contrast to commonly used ribosome profiling protocols, the use of cyclohexemide (CHX) was avoided, because CHX does not prevent disassembly of ribosomes located at TCs. Instead, cells were flash-frozen with liquid nitrogen, which has been shown to block elongating ribosomes as well as ribosomes located at TCs (Ingolia et al., 2011). Moreover, the use of CHX can bias the final output of ribosome footprints, as it seems to allow slow, concentration-dependent elongation prior to lysis Indeed according to recent studies (Gerashchenko and Gladyshev, 2014; Hussmann et al., 2015). The experiment was conducted in triplicates with ABCE1 KD and Control KD cells and the mapped reads showed overall strong 3 nucleotide periodicity, confirming that most of them indeed represent ribosome-protected RNA fragments (Fig. S5B). Ribosome occupancy was examined by dividing the transcripts into four regions (5’ UTR, CDS from the start codon to the middle, CDS from the middle to the TC, and 3’ UTR) and aligning ribosome-derived reads to each specific region, relative to the total number of reads (Fig. S5C). While ribosome occupancy in the 5’ UTR was low and very similar between ABCE1 KD and Control KD, a slightly lower ribosome coverage in the CDS was detected in ABCE1 KD compared to Control KD (Fig. S5C), indicating overall reduced translation and confirming the results of the puromycylation experiment (Fig. S4). On the contrary, on average more ribosomes appear to be present in 3’ UTR in the ABCE1 KD compared to Control KD, indicating that depletion of ABCE1 increases the rate of TC readthrough and/or translation re-initiation. These results are in agreement with other studies that have observed ribosome stalling at TCs and an increased ribosome occupancy in the 3’UTR upon ablation of ABCE1 (Mills et al., 2016; Sudmant et al., 2018; Young et al., 2015).

To obtain more detailed information about ribosome occupancy at TCs and in the 3’ UTR, we performed a metagene analysis of the ribosome-protected footprints in the range between 300 nucleotides upstream of the TC and 100 nucleotides downstream of the ABCE1 KD and the Control KD (Fig. 5A-B, top). While average ribosome occupancy at the TC seemed to be the same for ABCE1 KDs and Control KDs, an increased ribosome occupancy in the ABCE1 depleted cells was observed 30 nucleotides upstream of the TC and in the first 100 nucleotides of the 3’UTR (Fig. 5A-B, top), suggestive of disome accumulation at TCs and increased readthrough, respectively. As revealed by the heatmap, ribosome occupancy, and by inference translation of the ∼60’000 transcripts was affected differently by the depletion of ABCE1 (Fig. 5A, bottom). When ordered according to ribosome coverage at the TC in ABCE1 KD (Fig. 5A, bottom), about 20% of the transcripts exhibited marked ribosome stalling at the TC compared to Control KD (top section labelled by green line), while more than half of the transcripts showed decreased ribosome occupancy throughout the CDS and at the TC, gradually decreasing to a level at which no ribosomes were detected anymore at the TC (bottom section labelled by red line). The combination of these two effects explains why in the metagene analysis, no difference in ribosome occupancy at the TC was visible between ABCE1 KD and Control KD. By contrast, the increased ribosome occupancy in the 3’UTR in the metagene analysis was also clearly visible when individual transcripts were analyzed and ordered according to ribosome coverage in the 3’UTR in the ABCE1 KD dataset (Fig. 5B, bottom). Interestingly, of the 10’000 transcripts with the most ribosomes in the 3’ UTR (Fig. 5B, top section labelled by yellow line), one quarter also showed a concomitant higher ribosome coverage in the CDS. Therefore, to exclude that the increase of ribosome coverage in the 3’ UTR was simply due to a general high engagement of ribosomes with these specific transcripts, the ribosome-protected reads mapping in the 3’ UTR were normalized to the average ribosome density in the CDS (Fig. 5C). Confirming our previous notion, ABCE1 KD exhibited elevated ribosome occupancy in the 3’ UTR, even when scaled to ribosome occupancy in CDS, supporting the hypothesis that ABCE1 depletion might create an environment favourable for readthrough or re-intiation events. Moreover, the “bump” detected 30 nucleotides upstream of the TC in the trace of the metagene analysis specifically in the ABCE1 KD samples likely represents a second ribosome stalled immediately adjacent to ribosome located at the TC, since the distance from the TC matches with that recently reported for collided ribosomes (Ikeuchi et al., 2019). Hence, we propose that the high ribosome occupancy in CDS and 3’ UTR might represent ribosome stalling at the TC, which leads to ribosome queuing in front of the TC and occasional pushing of a ribosome past the TC, where it continues translation into the 3’ UTR. Our ribosome profiling data does not reveal if the ribosomes detected in the 3’ UTR are indeed still actively translating. However, TC readthrough has been reported previously as a mechanism that can suppress NMD (Hogg and Goff, 2010). Therefore, we checked where the previously identified endogenous NMD-sensitive transcripts (Colombo et al., 2017) were located in the heatmaps shown in Fig. 5A and B. When ribosome occupancy at the TC was considered, NMD targets were distributed homogeneously throughout the entire heatmap, being present in both the top (green) as well as the bottom (red) group (Fig. 5A). It therefore seems that ribosome stalling in the absence of ABCE1 occurs neither preferentially nor specifically on NMD targets, which might explain why the mRNA levels of only a fraction of NMD targets was upregulated in ABCE1 KD (Fig. 1B).

**Figure 5.**
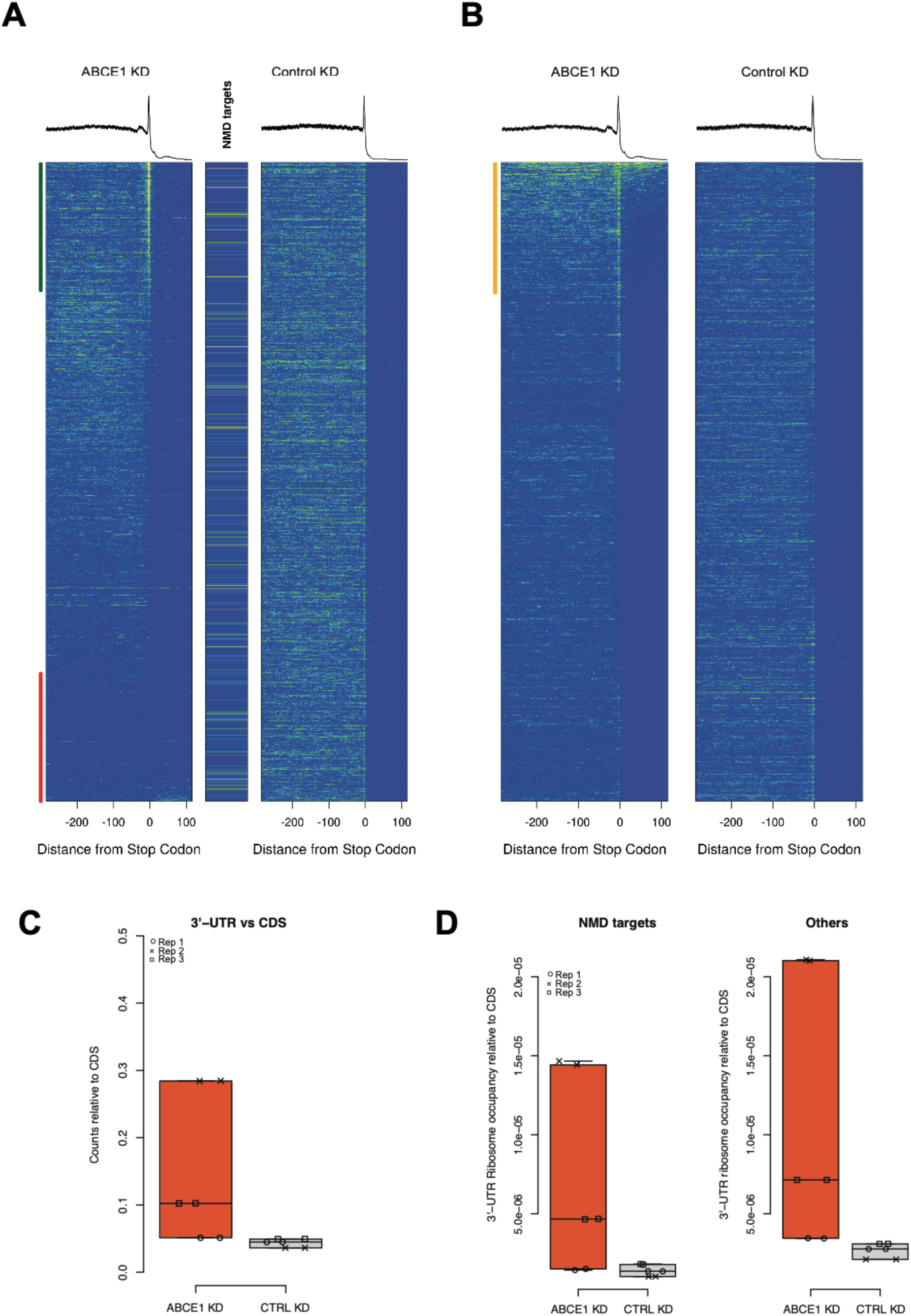
Enrichment of ribosome occupancy at termination codon and in the 3’ UTR in ABCE1 depleted cells. (A, top) Metagene analysis of ribosome-protected footprints from ABCE1 KD (left) or Control KD (right). Transcripts were aligned to the stop codon and mapped reads from 300 nucleotides upstream to 100 nucleotides downstream of the stop codon are shown. (A, bottom) Heatmap of ribosome-derived reads of all the transcripts considered for metagene analysis. Transcripts are ordered according to their ribosome occupancy at the termination codon in the ABCE1 KD samples. Transcripts at the top of the heatmap (top 10’000) with higher ribosome occupancy at the stop codon are marked with a green line. Transcripts at the bottom of the heatmap (bottom 10’000) with the lowest ribosome coverage at stop codon are marked with a red line. NMD sensitive transcripts identified in (Colombo et al., 2017) are shown between the heatmap panels. Each pixel corresponds to 10 transcripts. (B, top) Metagene analysis of ribosome coverage as in A, top. (B, bottom). Heatmap of ribosome-derived reads of all the transcripts considered for metagene analysis. Transcripts are ordered according to their ribosome occupancy in the 3’ UTR in the ABCE1 KD samples. Transcripts at the top of the heatmap (top 10’000) with high ribosome occupancy in the 3’ UTR are marked with an yellow line. (C) Analysis of ribosome occupancy in the 3’ UTR relative to mean ribosome occupancy in CDS. Total counts of ribosome-derived reads mapping in the 3’ UTR are plotted relative to total counts of ribosome-derived reads aligning to the CDS in each biological replicates for ABCE1 KD and Control KD. Percentage of reads relative to total reads are shown on the y-axis. (D) Analysis of ribosome occupancy in the 3’ UTR as in C, but performed separately for NMD targets and all other transcripts.

We next tried to search for those NMD targets upregulated in ABCE1 KD (identified in the RNA-seq experiment, Fig. 1B) in the green and red groups of transcripts, but because several of them share most of their sequence with the corresponding protein-coding isoform, unambiguous transcript identification was not possible and the analysis overall was inconclusive. However, since the NMD-sensitive SMG5-encoding mRNA consists of a single isoform, it was possible to unambiguously identify this transcript. It belongs to the top 10’000 transcripts with a high ribosome occupancy at the TC (Fig. 5A, section labelled by the green line). To still look at NMD targets separately and compare it to other transcripts that are insensitive to NMD, the ribosome occupancy in 3’ UTRs, relative to CDS, was analyzed as before (Fig. 5C), this time comparing the NMD targets against the rest of the transcripts (Fig. 5D). Generally, higher ribosome occupancy in the 3’ UTR was detected in both transcript categories. In NMD targets, translating ribosome in 3’ UTR can prevent NMD activation. Since ribosome profiling does not inform if the ribosome-protected RNA fragments detected in the 3’ UTR originate from continued translation after a readthrough event, from ribosomes that have re-initiated translation, or if they represent translationally inactive ribosomes without any peptide chain associated, we addressed this question in a reporter gene system.

### ABCE1 KD promotes readthrough on NMD sensitive transcripts

We noted that among the endogenous NMD targets (Colombo et al., 2017) that we detected in our RNA-seq, UGA was the most frequently occurring TC. Among these transcripts, the occurrence of the UGA TC was even further enriched among the sub-population of endogenous NMD targets that was upregulated upon ABCE1 KD: about 82% of them have UGA as the stop codon compared to 55% of the endogenous NMD targets that were not upregulated under ABCE1 depletion. According to previous studies, UGA is the “weakest” (i.e. least efficiently recognized) stop codon of the three canonical stop codons, resulting in the highest readthrough rates (Dabrowski et al., 2015). Because of the overrepresentation of the UGA stop codon among our ABCE1-dependent NMD targets, and since insertion of readthrough-promoting retroviral RNA elements into NMD reporter transcripts has been shown to antagonize UPF1 binding to their 3’ UTR and subsequent RNA decay (Hogg and Goff, 2010), we hypothesized that increased translational readthrough could suppress NMD on the NMD targets that were upregulated upon ABCE1 KD. To measure readthrough, we used a well-established readthrough reporter gene (Ivanov et al., 2008) in which the ORFs for Renilla luciferase (Rluc) and Firefly luciferase (Fluc) are positioned either in frame or separated by one of the three stop codons (Fig. 6A). The Fluc/Rluc ratio serves as a measure to quantify readthrough of stop codons under different conditions. First, we tested readthrough efficiency in cells transiently transfected with the dual-luciferase reporter and treated with geneticin (G418), an aminoglycoside antibiotic that promotes readthrough (Bukowy-Bieryllo et al., 2016). Consistent with previous reports (Loughran et al., 2014; Manuvakhova et al., 2000), basal readthrough frequency was lowest with UAA and somewhat higher with UAG and UGA. After treatment of the cells with 400 μg/mL G418 for 24 hours, readthrough increased at each of the three stop codons, with UGA being the most permissive, reaching about 2.5% readthrough (Fig. S6). When we measured readthrough occurrence under Control KD and ABCE1 KD conditions (Fig. 5B), we observed increased readthrough at all three stop codons upon ABCE1 depletion (Fig. 5C). ABCE1 KD caused the strongest increase in readthrough at the UAA stop codon, with UAA and UGA reaching 0.4% and 0.5% readthrough, respectively, while UAG was read through with about 0.2% efficiency. This result demonstrates that ABCE1 KD increases readthrough frequency in HeLa cells.

**Figure 6.**
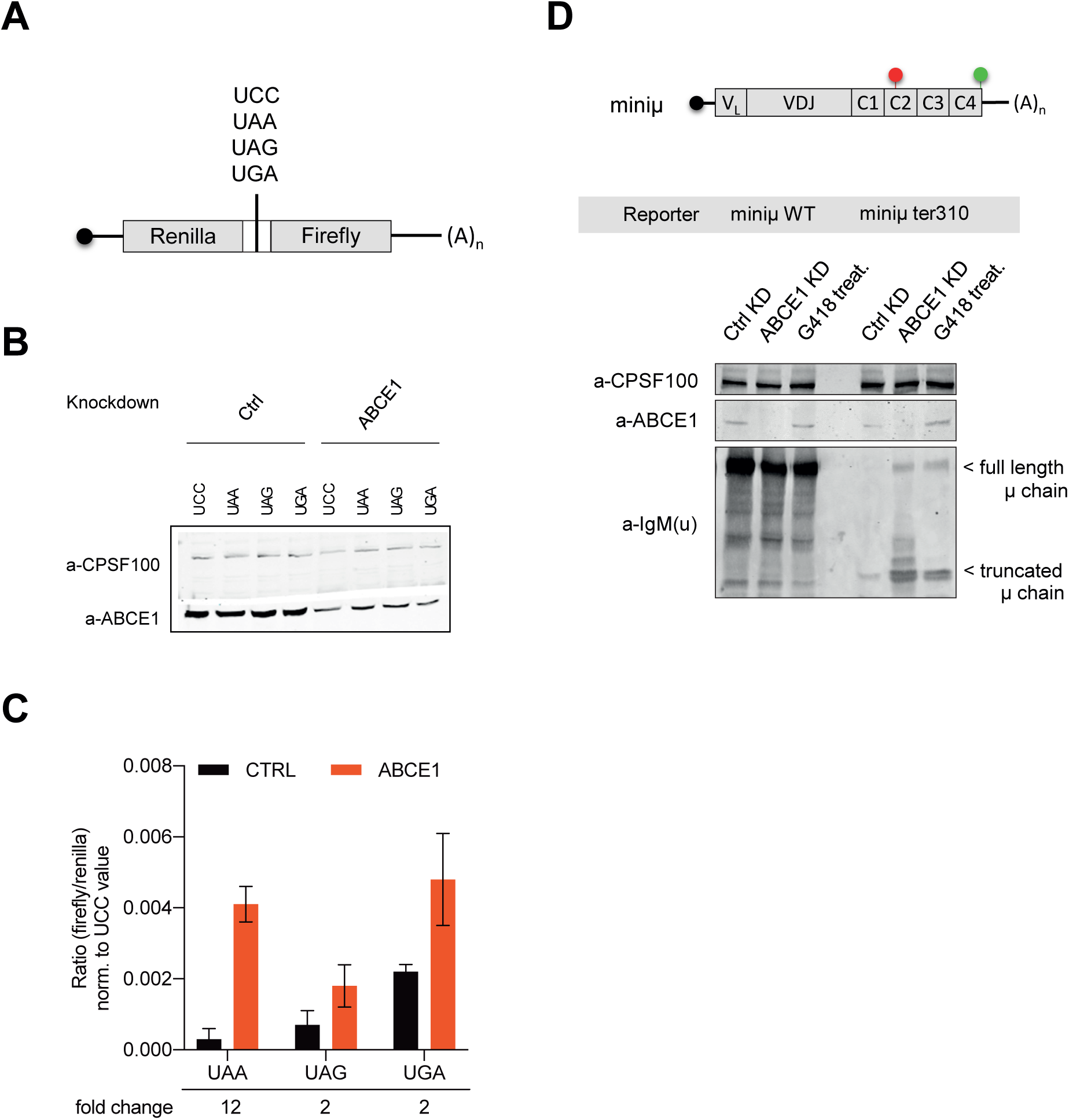
ABCE1 KD promotes readthrough on NMD sensitive transcripts. (A) Schematic representation of the dual luciferase readthrough reporter construct. (B) Protein lysates from HeLa cells transfected with siRNA against ABCE1 or a Control siRNA were immunoblotted with an anti-ABCE1 antibody. The blot is representative of at least three independently performed experiments with similar results. CPSF-100 served as loading control. (C) The ratios between firefly and renilla luciferase activities were determined for the stop codon-containing reporters and calculated relative to the non-stop reporter, representing the percentage of readthrough at the three different stop codons. HeLa cells depleted for ABCE1 (ABCE1 KD) or transfected with a control siRNA (CTRL) were co-transfected with dual luciferase constructs. Mean values and standard deviations from three independent experiments are shown. (D) Ig-μ protein level analysis by western blot of miniμ-expressing HeLa cells depleted of ABCE1 or after G418 treatment. Bands corresponding to truncated (translation termination at ter310) and full-length Ig-μ protein are indicated. CPSF-100 served as loading control. Efficiencies of ABCE1 depletion was also assessed. The blot is representative of at least three independently performed experiments with similar results.

To assess if readthrough might explain the observed NMD inhibition under ABCE1 KD, we examined if we could detect increased readthrough in NMD-sensitive transcripts. The Ig-μ minigene system provided an ideal opportunity for this, because of a polyclonal antibody that specifically recognizes the constant region of this mouse Ig-μ heavy chain. This antibody detects both the full-length Ig-μ chain as well as the truncated protein produced from the miniμ ter310. Western blotting was performed with lysates from cells stably expressing miniµ WT or ter310 mRNA and treated with control or ABCE1 siRNAs, or with G418 (Fig. 5D). While full-length Ig-μ was readily detected in comparable amounts in miniµ WT-expressing cells under all conditions, as expected, truncated Ig-μ chains were detected in cells expressing miniμ ter310. As a consequence of increased miniμ ter310 mRNA due to NMD inhibition upon ABCE1 KD or G418 treatment, the amount of truncated Ig-μ chains increased. Most importantly, in miniμ ter310-expressing cells depleted for ABCE1 or treated with G418, full-length Ig-μ chains were also detected in addition to the truncated ones, indicating that under these conditions, a sizable fraction of the ribosomes read through the PTC and translated all the way down to the normal termination codon, thereby presumably displacing UPF1, EJCs and other NMD-inducing factors from the region downstream of the PTC. Noteworthy, the detection of full-length Ig-μ chains in miniμ ter310-expressing ABCE1-depleted cells suggests that ABCE1 not only functions in splitting and recycling of the small and large ribosomal subunits, but that it also plays a role in polypeptide release during translation termination.

## DISCUSSION

The ribosome recycling factor ABCE1 has so far been shown to be required for splitting apart the large and small ribosomal subunits after translation termination (Gerovac and Tampe, 2019) and under conditions of limiting amounts of eRF3 to stimulate eRF1-mediated peptide release (Shoemaker and Green, 2011). Here we show that, somewhat unexpectedly, ABCE1 is also needed for correct activation of NMD, suggesting that correct recycling of ribosomes is a pre-requisite for NMD. We report that depletion of ABCE1 suppresses NMD of a subgroup of NMD sensitive mRNAs in HeLa cells (Fig. 1) and of two well-established NMD reporters (i.e. TCRβ and miniμ) (Fig. 2). According to a previous study, the conditional depletion of the yeast orthologue of ABCE1, Rli1 causes ribosome stalling at the termination codon (TC), blocking degradation on NMD targets and leading to the stabilization of degradation-intermediates for NMD sensitive mRNAs (Serdar et al., 2016). Although NMD shares similarities between yeast and human cells, we only detected an increase of full-length RNA with the two NMD reporter genes TCRβ and miniμ under ABCE1 KD (Fig. 3), indicating that ABCE depletion inhibits NMD at a step prior to SMG6-mediated endonucleolytic cleavage. Since ABCE1 has been reported to function in translation termination and in the first steps of translation initiation (Gerovac and Tampe, 2019), its depletion is expected to affect overall protein synthesis, which in turn might cause the observed NMD inhibition, since NMD is strictly translation-dependent (Karousis and Muhlemann, 2019). However, although overall protein synthesis was reduced by about 40% in ABCE1 depleted cells (Fig. S4), this cannot account for the observed NMD inhibition. First, under ABCE1 KD conditions, NMD sensitive mRNAs were found strongly enriched in polysomes (Fig. 4), and secondly, protein products were detected for the miniµ reporter gene (Fig. 6D), excluding the possibility that NMD inhibition induced by ABCE1 depletion might simply result from an overall translation inhibition.

Readthrough has been previously reported to antagonize NMD by displacing UPF1 from the 3’ UTR of NMD reporter transcripts (Hogg and Goff, 2010). While rare readthrough events still allowed UPF1 re-association with the RNA, frequent readthrough led to a marked decrease of steady-state UPF1 interaction with the 3’ UTR and inhibited NMD (Hogg and Goff, 2010). If the mere clearing of NMD factors from the 3’ UTR by elongating ribosomes was sufficient to spare an mRNA from NMD, this could happen both by readthrough of the TC as well as by re-initiation of post-termination ribosomes downstream of the TC. Interestingly, in the absence of Rli1/ABCE1, an increase of ribosome occupancy in the 3’ UTR and re-initiation events have been reported in yeast and in anucleate platelets and reticulocytes (Mills et al., 2016; Young et al., 2015). Our results corroborate the idea of readthrough or re-initiation leading to NMD inhibition, as our ribosome profiling experiments detected increased ribosome occupancy in the 3’ UTR of many mRNAs (Fig. 5A-C), including many NMD sensitive mRNAs (Fig. 5D) in ABCE1 depleted cells. Indeed, full-length protein was detected from the PTC-harboring miniμ mRNA only in cells depleted for ABCE1 or when treated with a readthrough-promoting drug (G418), demonstrating that a fraction of the stalled ribosomes read through the PTC without releasing the nascent polypeptide and continue translation all the way to the normal TC (Fig. 6D). While for the miniμ NMD reporter, these readthrough events can explain the NMD inhibition, one has to postulate a slightly different mechanism to explain how the endogenous NMD targets that are upregulated in ABCE1 KD escape NMD, because the majority of them has several in-frame TCs closely downstream of the NMD-triggering TC, which would quickly stop again “escaping” ribosomes and prevent them from translocating some distance into the 3’ UTR and so from clearing the 3’ UTR of NMD factors. However, in most of the cases, one of the other two frames (the +1 or +2 frame) would allow translation to proceed past the last exon-exon junction, where termination is predicted to no longer trigger NMD. We therefore postulate that the readthrough events promoted by ABCE1 depletion frequently are accompanied by a frameshift of the ribosome. If indeed the readthrough of the TC is caused by a ribosome collision pushing the stalled ribosome at the TC downwards (see below), it seems plausible that the pushed forward ribosome might resume translation randomly in any of the three reading frames. It remains to be tested whether such frameshifting readthrough indeed occurs in the absence of ABCE1.

It is currently not understood how ABCE1 depletion leads to readthrough and why the nascent polypeptide does not get efficiently released. According to our current understanding of translation termination, ribosome recycling at the TC is expected occur after polypeptide release, preventing readthrough and translation into the 3’ UTR (Hellen, 2018). Yet our ribosome profiling results showed clearly an increase of ribosome occupancy at the TC in ABCE1 KD cells, as well as an enrichment of ribosome coverage in the 3’ UTR (Fig. 5A and B), consistent with previous studies (Mills et al., 2016; Sudmant et al., 2018; Young et al., 2015). However, not only termination seems to be affected in the absence of ABCE1, as about the 20% of all transcripts analyzed in the ribosome profiling do not show any ribosomes at the TC and in the CDS (Fig. 5A, section marked by red line), suggesting that translation of these specific transcripts ceased when ABCE1 became limiting. In contrast, transcripts with a high ribosome occupancy in the 3’ UTR tend to have more ribosomes also in the CDS (this applies to 6% of all analyzed transcripts). Interestingly, the metagene analysis of ribosome occupancy revealed an accumulation of ribosome-protected reads 30 nucleotides upstream the TC exclusively in the ABCE1 depleted cells (Fig. 5A and B), which most likely represents ribosomes sitting immediately behind the ribosome stalled at the TC. This signal at −30 nucleotides is also clearly visible by eye in the heatmap on the fraction of transcripts that exhibits the strongest ribosome stalling at the TC (Fig. 5A, top section labelled by the green line). This further contributes to the idea of a ribosome traffic jam on specific mRNAs under ABCE1 depletion as a consequence of improper ribosome clearing at the TC. This prolonged stalling at the TC increases the chance for a ribosome to read through the TC or to re-initiate translation downstream of the TC if the nascent peptide has already been released. Since the transcripts showing a −30 nucleotide “bump” do not all have increased ribosome occupancy in the 3’ UTR, it seems that we have captured two different outcomes of ABCE1 KD: on a fraction of the transcripts, ribosomes accumulate immediately upstream of the ribosome stalled at the TC, while on another transcript population, ribosomes strongly accumulate in the CDS, at TC and in the 3’ UTR. While on the former group of transcripts, it seems that the ribosomes are stuck at the stop codon but translation rate on these transcripts is low, which prevents the pileup of ribosomes along the CDS, the latter group might represent a traffic jam of ribosomes that accumulate on heavily translated transcripts because of delayed release at the TC. Our hypothesis is that some of the stalled ribosomes at the TC might read through the TC by incorporating a near-cognate tRNA, whereas others could be “forced” to advance beyond the mRNA, pushed by a following ribosome running into the on stalled at the TC. It seems plausible that in the second scenario, the forward pushed ribosome could resume translation in another frame, which would explain for many of the endogenous NMD targets that are upregulated in the ABCE1 KD how their 3’ UTR can be cleared of NMD-inducing factors, in particular from UPF1 and EJCs. Moreover, the traffic jam model might also explain the heterogeneous ribosome distribution across all transcripts, with a group of transcripts being highly translated and a fraction of mRNAs without any ribosome associated, since the highly-translated transcripts might sequester a sizable fraction of the ribosomes, thereby preventing their recycling and association with other mRNAs. It was reported that the intracellular abundance of the release factor eRF1 plays a crucial role in TC recognition and readthrough (Cridge et al., 2018), and limiting eRF1 availability by trapping it ribosomes stalled at TCs on a subset of mRNAs could lead to enhanced readthrough on other transcripts. The fact that we observed readthrough without peptide release – i.e. full-length protein production – under ABCE1 KD could also be the result of such a sequestering of eRF1.

Further investigations are ongoing to gain more insight on the translation level in cells depleted for ABCE1 and to clarify the traffic jam theory and the sponge hypothesis. The recently developed method that allows measuring translation termination events in real-time on single mRNA molecules (Yan et al., 2016) is a powerful method that will shed light on the translation and translation termination dynamics of NMD sensitive and insensitive mRNAs under ABCE1 KD. Additionally, a detailed study to clarify re-initiation and readthrough rate on NMD sensitive mRNAs should also be considered in the future, using a reporter designed to discriminate between the two mechanisms. Certainly, ABCE1 plays a general role in translation and, as reported also in this study, its depletion did not specifically affect only NMD. However, deciphering the translation termination mechanism and its impact on NMD targets is an approach that can help understanding how NMD activation is directly connected to translation termination and identifying factors that promote or inhibit proper termination, thereby controlling NMD activation.

## MATERIAL AND METHODS

### Cell culture and transfection

Hela cells were cultured in DMEM supplemented with 10% fetal calf serum (FCS) and antibiotics. Cells were grown in 5% CO_2_ at 37°C. HeLa cells stably expressing TCRβ and the Ig-μ reporter genes were described previously (Buhler et al., 2004; Mohn et al., 2005). For readthrough experiments, cells were treated with 400 ug/mL G418 (Fisher Scientific) for 24 hours before harvesting. SiRNA-mediated knockdowns were carried out by a double-transfection procedure. First, 3*10^5^ HeLa cells per well were seeded in 6-well plates. One day later, the cells were transfected with 52 pmol of siRNA using Lullaby reagent (OZ Biosciences). After 2 days, the cells were re-transfected as before. Protein and total RNA were isolated after one additional day. SiRNAs with the sequence 5’-GAGGAGAGUUGCAGAGAUUU-3’ for targeting ABCE1 and 5’-GAUGCAGUUCCGCUCCAUU-3’ for targeting UPF1 were used. Readthrough assays were performed using the p2luc plasmids as described previously (Ivanov et al., 2008). Briefly, 3*10^5^ HeLa cells per well were seeded in 6-well plates. One day later, the cells were transfected with 52 pmol of siRNA using Lullaby reagent (OZ Biosciences). After 2 days, the cells were re-transfected with 52 pmol of siRNA and 800 ng of the indicated plasmids using Lipofectamine 2000 (Invitrogen). Protein and total RNA were isolated after two additional days. Knockdown efficiencies of the different targets were verified by western blotting using anti-RENT1 (UPF1) (Bethyl, A300–038A), anti-ABCE1 (Abcam, ab185548), anti-Tyrosine Tubulin (Tub1A2) (Sigma, T9028) and anti-CPSF100 (custom made) antibodies. For IgM-µ protein detection, anti-IgM-µ (Jackson Imm. 115-005-020) was used in combination with Pierce Western Blot Signal Enhancer kit (Thermo Scientific).

### Luciferase assay for readthrough quantification

Dual luciferase assays were performed according to the manufacturer’s protocol (Promega). Luminescence was measured using TECAN Infinite M1000, two technical replicates for each sample. Readthrough efficiency was calculated by comparing the ratio of firefly to renilla luciferase activity relative to the plasmid without stop codon.

### Puromycin incorporation assay

Translation level analysis was performed using an adapted protocol from a previous study (Schmidt 2009). Briefly, HeLa cells were labeled with 10 μg/ml of puromycin (Santa Cruz) for 10 min in complete medium. After 30 minutes in puromycin free medium, cells were harvested and puromycin incorporation was detected by immunoblotting using a specific anti-Puromycin-12D10 antibody (Millipore, MABE343). The western blot signal was quantified using the ImageJ program.

### RNA manipulation and analysis

RNA was extracted from cells using TriReagent according to the manufacturer’s instructions. RT-qPCR analyses were performed using Brilliant III Ultra-Fast SYBR® Green (Agilent) after DNase treatment using Turbo DNase (AMBION). All primers used in this study are listed in Supplementary Table 1. For the analysis of total RNA by RNA-seq, rRNA-depleted mRNA was purified and used for library construction using TruSeq Stranded mRNA Library Prep Kit (Illumina) and sequenced with Illumina HiSeq2500. Northern blot analysis was performed according to an adapted protocol from a previous study (Eberle et al., 2009). Briefly, approximately 10 µg RNA per sample was separated on a 1,2% (w/v) agarose gel containing 1x MOPS and 1% (v/v) formaldehyde and transferred to a positively charged nylon membrane by wet transferring. The ^32^P-labeled ribo-probe was hybridized in ULTRAHyb buffer (AMBION) at 68°C overnight. The ribo-probes were transcribed from a NotI digested plasmids using SP6 polymerase (Fisher) in the presence of α^32^P-UTP. The ribo-probe sequences are listed in Supplementary Table 1.

### Polysome fractionation

The polysome profiling protocol was adapted from (Zuccotti and Modelska, 2016). Briefly, HeLa cells, with or without knockdown (two 150-mm culture dishes each condition) were treated with 100 µg/ml cycloheximide (Focus biomolecules) for 4 min at 37 °C. Cells were then lysed in low-salt buffer (10 mM Tris-HCl [pH 7.5], 10 mM NaCl, 10 mM MgCl_2_, 1 mM DTT, 1% Triton X-100, 1% Na-Deoxycholate) supplemented with RNAse Inhibitor (NxGen), Protease Inhibitor cocktail (Biotools) and 100 μg/ml cycloheximide by quickly vortexing. After 2 min on ice, lysates were cleared by centrifugation for 5 min at 16’000 g. Lysates were loaded on 15–50% sucrose gradient tube and centrifuged at 40’000 rpm in a Beckman SW-41Ti rotor for 2 h at 4 °C. Gradients were fractionated and monitored at absorbance 254 nm with a fraction collector (BioComp Instruments) at a 0.2-mm/s EM1 speed, with a distance of 3.71 mm per fraction. About two hundred microliters of each fraction was used for RNA extraction using ethanol precipitation followed by TriReagent procedure.

### Ribosome profiling experiment

Cells for ribosome profiling and RNA sequencing were processed using an adapted protocol from previous publications (Aeschimann et al., 2015; Ingolia et al., 2012). Cells were washed with ice-cold phosphate buffered saline (PBS) and flash-frozen in liquid nitrogen. Subsequently, cells were scraped and lysed in lysis buffer (20 mM Tris HCL pH7.4, 150 mM NaCl, 5 mM MgCl_2_, 1% Triton X-100, 1 mM DTT, 25 U/uL Turbo DNase, Turbo DNase buffer) on ice. Next, cells were triturated ten times through a 27-gauge needle of a syringe and clarified by centrifugation. For ribosome profiling, lysates (about 4 U_260_) were subsequently treated with 200 U RNase I (Ambion) for 10 min at 23°C and shaking at 300 rpm. The digestion was stopped by addition of 100 U SUPERase In RNase inhibitor (Ambion). Monosomes were separated on Illustra Micro-Spin S-400 HR gel filtration columns (GE Healthcare Life Science) as previously described (Aeschimann et al., 2015). TriReagent was added immediately to the eluates and samples were stored at −80°C until further processing. RNA was isolated according to the TriReagent protocol and separated on 15% Novex polyacrylamide gels (Invitrogen). Ribosome footprints were excised between 26 and 34 nucleotide RNA size markers. After RNA isolation and purification, rRNAs were removed using the RiboZero kit (Illumina) according to manufacturer’s datasheet. Sequencing libraries from ribosome footprints were generated as previously described (Aeschimann et al., 2015). RNA-seq libraries were prepared from cells lysed similar as the ribosome profiling samples. RNA was isolated from cleared lysates by addition of TriReagent as for the ribosome-protected fragments (see above). Total RNA was used for library generation with the TruSeq Stranded mRNA Library Prep Kit (Illumina) according to the manufacturer’s instructions. Libraries were sequenced on an Illumina HiSeq2500 generating 100 nt single-end reads.

## ACKNOWLEDGMENTS

We are grateful to Martinio Colombo for initial bioinformatics analyses and thank Evan Karousis, Andrea Eberle, Sofia Nasif, Lara Contu and Lukas Gurzeler for thoughtful comments. This work was supported by the NCCR RNA & Disease funded by the Swiss National Science Foundation (SNSF), by SNSF grants 31003A-162986 and 310030B-182831, and by the canton of Bern.

## AUTHOR CONTRIBUTIONS

G.A. and O.M designed the study and planned the experiments; G.A. and M.D. performed all wetlab experiments; R.D. and S.C. performed bioinformatics analyses; G.A. and O.M. wrote the paper with inputs from M.D., R.D. and S.C.

## SUPPLEMENTAL INFORMATION

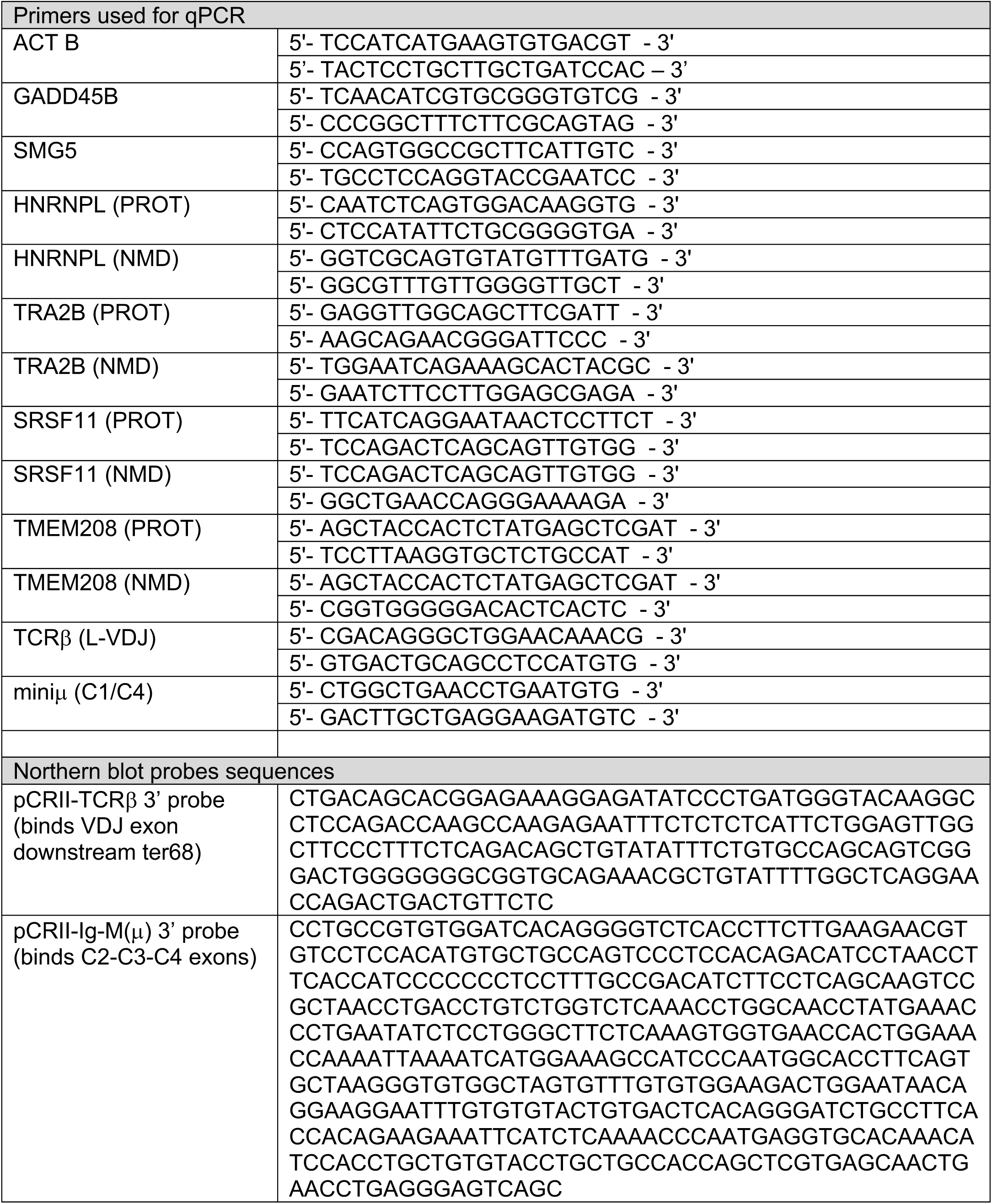

**Figure S1.**
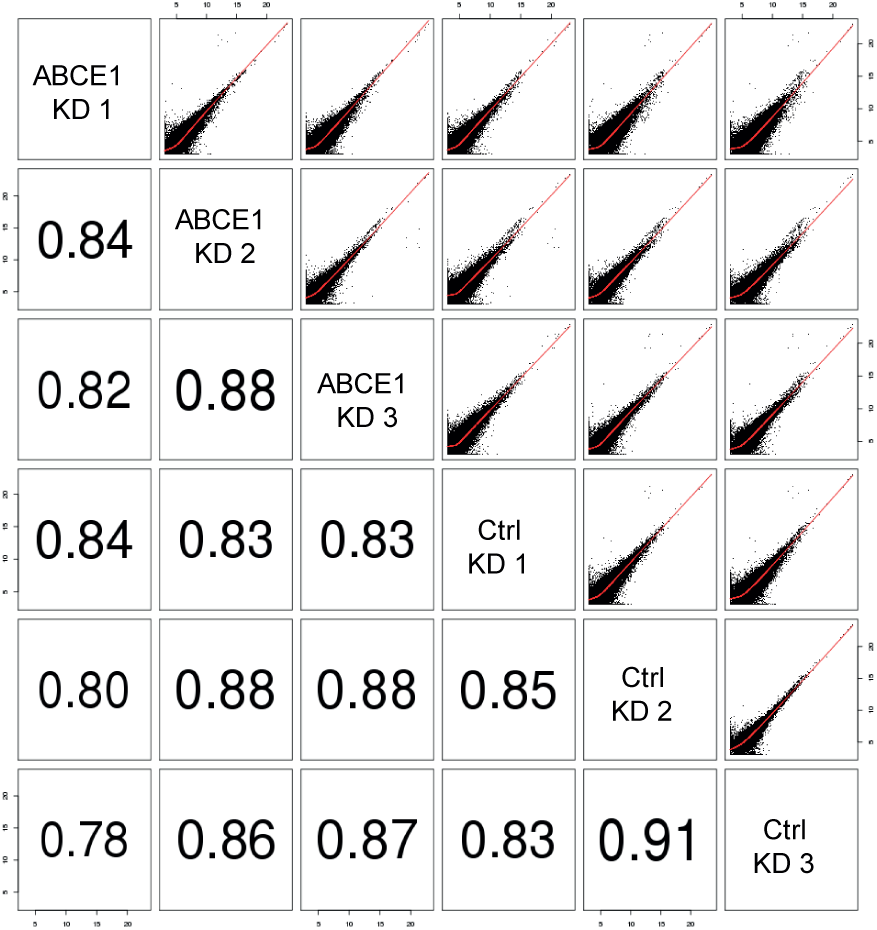
mRNA expression in ABCE1 KD cells. Correlation of mRNA expression levels between three biological replicates in Control KD (Ctrl) and ABCE1 KD in RNA sequencing results. Pearson correlation coefficients are shown.

**Figure S4.**
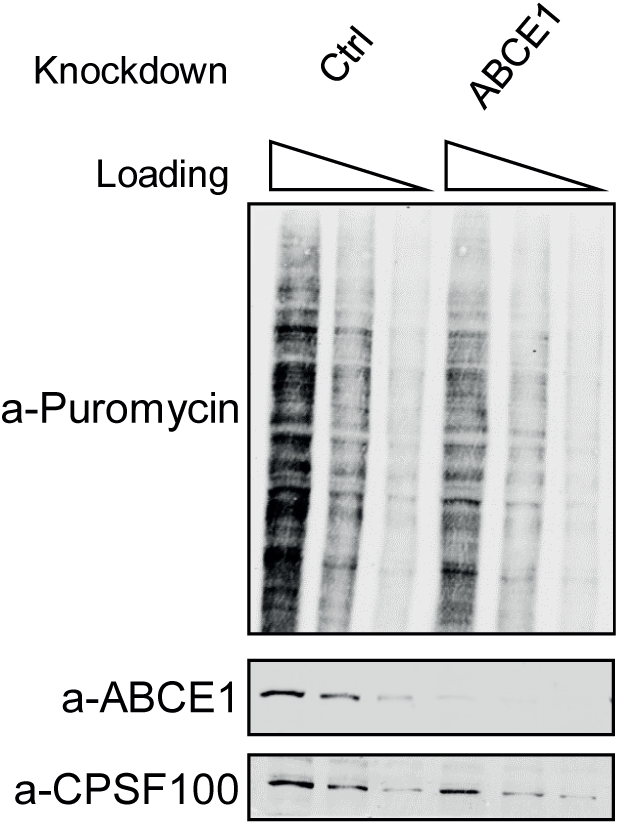
ABCE1 depletion reduces general translation. Translation levels were measured using SUnSET technique (Schmidt et al., 2009). Western blot analysis of HeLa cells under ABCE1 KD, labeled with puromycin and then analyzed with antibody against puromycin. ACTB immunoblot is shown as a loading control. Efficiencies of ABCE1 depletion is also measured. The blot is representative of at least three independently performed experiments with similar results. Three different amounts of Control KD and ABCE1 KD were loaded (100%, 50%, 25%).

**Figure S5.**
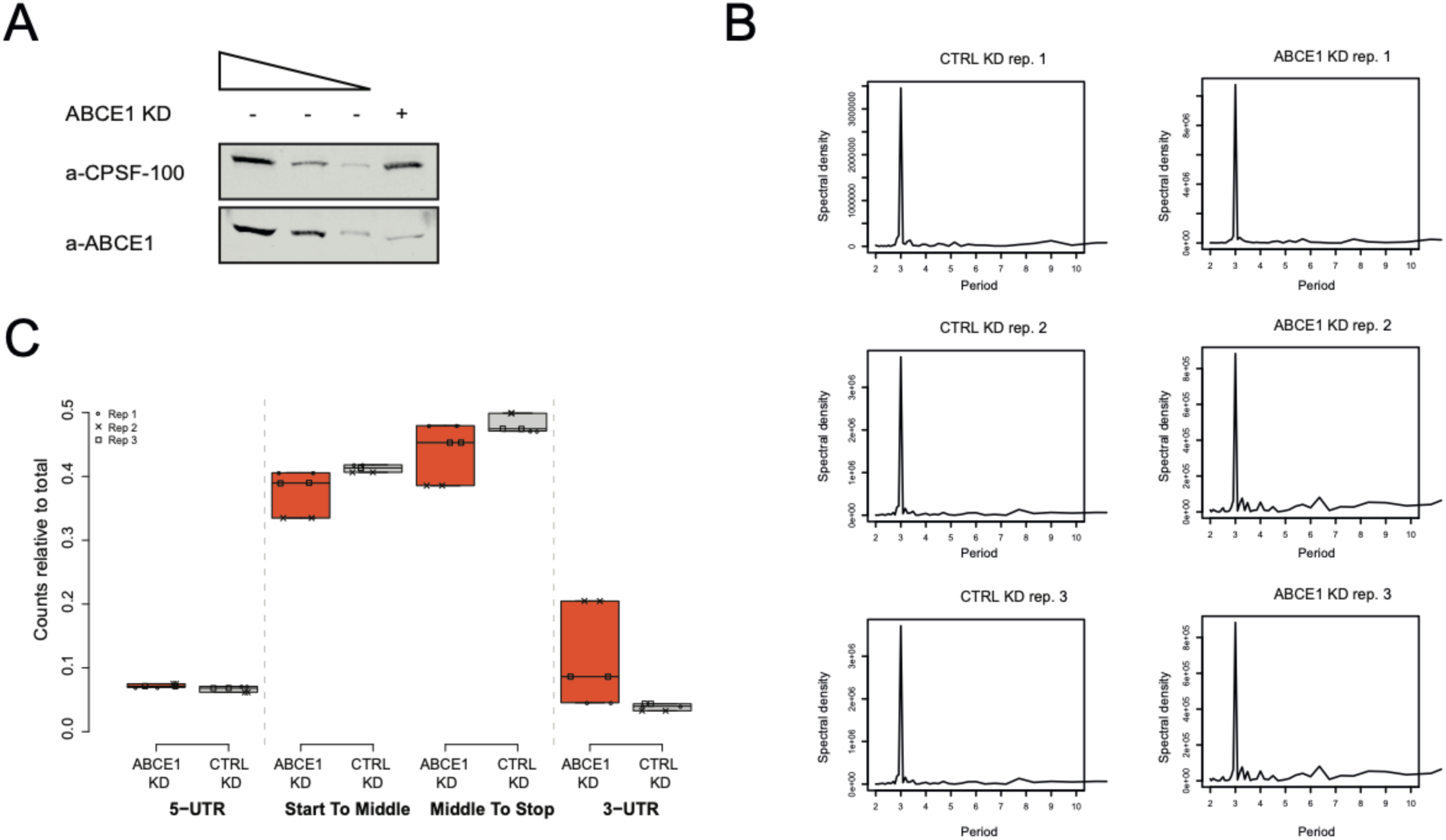
Ribosome profiling in ABCE1 depleted cells. (A) Protein level analysis by western blot of ABCE1 KD in ribosome profiling samples. CPSF-100 serves as loading control. Three different amounts of Control KD are loaded (100%, 30%, 10%). Thre blot is representative of the three independently performed biological replicates. (B) Reads aligned to the CDS start have been analysed. Fourier transformation analysis was applied to quantify the strength of the periodicity signal. This function returns the strength (spectral density) of each frequency found in periodic signals. A peak at period 3 indicates that reads distribution have 3 nucleotides periodicity. All the three replicates for each condition are shown. (C) Ribosome-derived reads aligned to different transcript regions. Total counts of ribosome-derived reads mapping in each region are plotted. Each biological replicate for ABCE1 KD and Control KD is shown. Percentage of reads relative to total reads are depicted on the y-axis.

**Figure S6.**
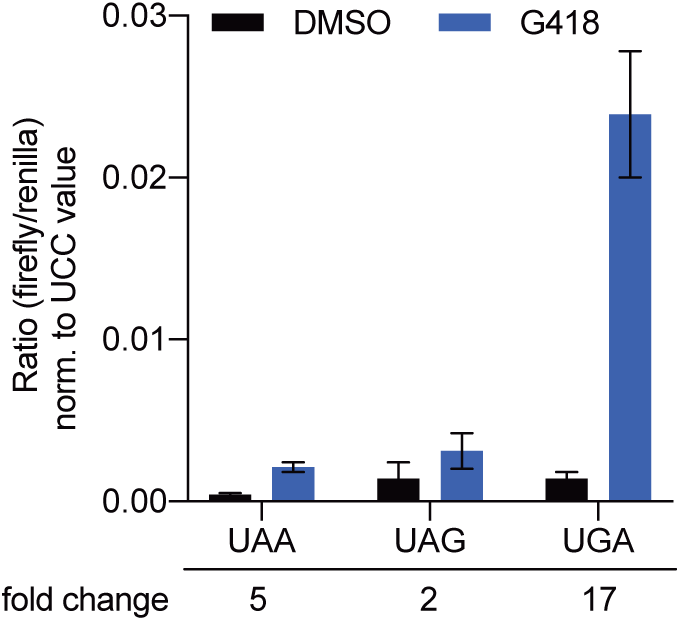
Positive control for readthrough assay. The blot shows the ratios between firefly and renilla luciferase activities, representing the percentage of readthrough at the three different stop codons. HeLa cells were transfected with dual luciferase constructs and then treated for 24 hours with G418. The firefly:renilla ratio obtained with the non-stop construct was set as 100% readthrough. Mean values and standard deviations from three independent experiments were calculated.

## REFERENCES

Aeschimann, F., Xiong, J., Arnold, A., Dieterich, C., and Grosshans, H. (2015). Transcriptome-wide measurement of ribosomal occupancy by ribosome profiling. Methods 85, 75–89.

Amrani, N., Ganesan, R., Kervestin, S., Mangus, D.A., Ghosh, S., and Jacobson, A. (2004). A faux 3’-UTR promotes aberrant termination and triggers nonsense-mediated mRNA decay. Nature 432, 112–118.

Behm-Ansmant, I., Gatfield, D., Rehwinkel, J., Hilgers, V., and Izaurralde, E. (2007). A conserved role for cytoplasmic poly(A)-binding protein 1 (PABPC1) in nonsense-mediated mRNA decay. EMBO J 26, 1591–1601.

Behm-Ansmant, I., and Izaurralde, E. (2006). Quality control of gene expression: a stepwise assembly pathway for the surveillance complex that triggers nonsense-mediated mRNA decay. Genes Dev 20, 391–398.

Buhler, M., Paillusson, A., and Muhlemann, O. (2004). Efficient downregulation of immunoglobulin mu mRNA with premature translation-termination codons requires the 5’-half of the VDJ exon. Nucleic Acids Res 32, 3304–3315.

Buhler, M., Steiner, S., Mohn, F., Paillusson, A., and Muhlemann, O. (2006). EJC-independent degradation of nonsense immunoglobulin-mu mRNA depends on 3’ UTR length. Nat Struct Mol Biol 13, 462–464.

Bukowy-Bieryllo, Z., Dabrowski, M., Witt, M., and Zietkiewicz, E. (2016). Aminoglycoside-stimulated readthrough of premature termination codons in selected genes involved in primary ciliary dyskinesia. RNA Biol 13, 1041–1050.

Calviello, L., Mukherjee, N., Wyler, E., Zauber, H., Hirsekorn, A., Selbach, M., Landthaler, M., Obermayer, B., and Ohler, U. (2016). Detecting actively translated open reading frames in ribosome profiling data. Nat Methods 13, 165–170.

Carlevaro-Fita, J., Rahim, A., Guigo, R., Vardy, L.A., and Johnson, R. (2016). Cytoplasmic long noncoding RNAs are frequently bound to and degraded at ribosomes in human cells. RNA 22, 867–882.

Carter, M.S., Doskow, J., Morris, P., Li, S., Nhim, R.P., Sandstedt, S., and Wilkinson, M.F. (1995). A regulatory mechanism that detects premature nonsense codons in T-cell receptor transcripts in vivo is reversed by protein synthesis inhibitors in vitro. J Biol Chem 270, 28995–29003.

Cho, H., Han, S., Choe, J., Park, S.G., Choi, S.S., and Kim, Y.K. (2013). SMG5-PNRC2 is functionally dominant compared with SMG5-SMG7 in mammalian nonsense-mediated mRNA decay. Nucleic Acids Res 41, 1319–1328.

Colombo, M., Karousis, E.D., Bourquin, J., Bruggmann, R., and Muhlemann, O. (2017). Transcriptome-wide identification of NMD-targeted human mRNAs reveals extensive redundancy between SMG6- and SMG7-mediated degradation pathways. RNA 23, 189–201.

Cridge, A.G., Crowe-McAuliffe, C., Mathew, S.F., and Tate, W.P. (2018). Eukaryotic translational termination efficiency is influenced by the 3’ nucleotides within the ribosomal mRNA channel. Nucleic Acids Res 46, 1927–1944.

Dabrowski, M., Bukowy-Bieryllo, Z., and Zietkiewicz, E. (2015). Translational readthrough potential of natural termination codons in eucaryotes--The impact of RNA sequence. RNA Biol 12, 950–958.

Eberle, A.B., Lykke-Andersen, S., Muhlemann, O., and Jensen, T.H. (2009). SMG6 promotes endonucleolytic cleavage of nonsense mRNA in human cells. Nat Struct Mol Biol 16, 49–55.

Eberle, A.B., Stalder, L., Mathys, H., Orozco, R.Z., and Muhlemann, O. (2008). Posttranscriptional gene regulation by spatial rearrangement of the 3’ untranslated region. PLoS Biol 6, e92.

Gehring, N.H., Lamprinaki, S., Kulozik, A.E., and Hentze, M.W. (2009). Disassembly of exon junction complexes by PYM. Cell 137, 536–548.

Gerashchenko, M.V., and Gladyshev, V.N. (2014). Translation inhibitors cause abnormalities in ribosome profiling experiments. Nucleic Acids Res 42, e134.

Gerovac, M., and Tampe, R. (2019). Control of mRNA Translation by Versatile ATP-Driven Machines. Trends Biochem Sci 44, 167–180.

Guydosh, N.R., and Green, R. (2014). Dom34 rescues ribosomes in 3’ untranslated regions. Cell 156, 950–962.

Hellen, C.U.T. (2018). Translation Termination and Ribosome Recycling in Eukaryotes. Cold Spring Harb Perspect Biol 10.

Heuer, A., Gerovac, M., Schmidt, C., Trowitzsch, S., Preis, A., Kotter, P., Berninghausen, O., Becker, T., Beckmann, R., and Tampe, R. (2017). Structure of the 40S-ABCE1 post-splitting complex in ribosome recycling and translation initiation. Nat Struct Mol Biol 24, 453–460.

Hogg, J.R., and Goff, S.P. (2010). Upf1 senses 3’UTR length to potentiate mRNA decay. Cell 143, 379–389.

Hurt, J.A., Robertson, A.D., and Burge, C.B. (2013). Global analyses of UPF1 binding and function reveal expanded scope of nonsense-mediated mRNA decay. Genome Res 23, 1636–1650.

Hussmann, J.A., Patchett, S., Johnson, A., Sawyer, S., and Press, W.H. (2015). Understanding Biases in Ribosome Profiling Experiments Reveals Signatures of Translation Dynamics in Yeast. PLoS Genet 11, e1005732.

Ikeuchi, K., Tesina, P., Matsuo, Y., Sugiyama, T., Cheng, J., Saeki, Y., Tanaka, K., Becker, T., Beckmann, R., and Inada, T. (2019). Collided ribosomes form a unique structural interface to induce Hel2-driven quality control pathways. EMBO J 38.

Ingolia, N.T., Brar, G.A., Rouskin, S., McGeachy, A.M., and Weissman, J.S. (2012). The ribosome profiling strategy for monitoring translation in vivo by deep sequencing of ribosome-protected mRNA fragments. Nat Protoc 7, 1534–1550.

Ingolia, N.T., Lareau, L.F., and Weissman, J.S. (2011). Ribosome profiling of mouse embryonic stem cells reveals the complexity and dynamics of mammalian proteomes. Cell 147, 789–802.

Ivanov, P.V., Gehring, N.H., Kunz, J.B., Hentze, M.W., and Kulozik, A.E. (2008). Interactions between UPF1, eRFs, PABP and the exon junction complex suggest an integrated model for mammalian NMD pathways. EMBO J 27, 736–747.

Karousis, E.D., and Muhlemann, O. (2019). Nonsense-Mediated mRNA Decay Begins Where Translation Ends. Cold Spring Harb Perspect Biol 11.

Kashima, I., Yamashita, A., Izumi, N., Kataoka, N., Morishita, R., Hoshino, S., Ohno, M., Dreyfuss, G., and Ohno, S. (2006). Binding of a novel SMG-1-Upf1-eRF1-eRF3 complex (SURF) to the exon junction complex triggers Upf1 phosphorylation and nonsense-mediated mRNA decay. Genes Dev 20, 355–367.

Keeling, K.M., Lanier, J., Du, M., Salas-Marco, J., Gao, L., Kaenjak-Angeletti, A., and Bedwell, D.M. (2004). Leaky termination at premature stop codons antagonizes nonsense-mediated mRNA decay in S. cerevisiae. Rna 10, 691–703.

Le Hir, H., Gatfield, D., Izaurralde, E., and Moore, M.J. (2001). The exon-exon junction complex provides a binding platform for factors involved in mRNA export and nonsense-mediated mRNA decay. EMBO J 20, 4987–4997.

Le Hir, H., Izaurralde, E., Maquat, L.E., and Moore, M.J. (2000). The spliceosome deposits multiple proteins 20-24 nucleotides upstream of mRNA exon-exon junctions. EMBO J 19, 6860–6869.

Li, Z., Vuong, J.K., Zhang, M., Stork, C., and Zheng, S. (2017). Inhibition of nonsense-mediated RNA decay by ER stress. RNA 23, 378–394.

Loh, B., Jonas, S., and Izaurralde, E. (2013). The SMG5-SMG7 heterodimer directly recruits the CCR4-NOT deadenylase complex to mRNAs containing nonsense codons via interaction with POP2. Genes Dev 27, 2125–2138.

Loughran, G., Chou, M.Y., Ivanov, I.P., Jungreis, I., Kellis, M., Kiran, A.M., Baranov, P.V., and Atkins, J.F. (2014). Evidence of efficient stop codon readthrough in four mammalian genes. Nucleic Acids Res 42, 8928–8938.

Lykke-Andersen, J., and Bennett, E.J. (2014). Protecting the proteome: Eukaryotic cotranslational quality control pathways. J Cell Biol 204, 467–476.

Mancera-Martinez, E., Brito Querido, J., Valasek, L.S., Simonetti, A., and Hashem, Y. (2017). ABCE1: A special factor that orchestrates translation at the crossroad between recycling and initiation. RNA Biol 14, 1279–1285.

Manuvakhova, M., Keeling, K., and Bedwell, D.M. (2000). Aminoglycoside antibiotics mediate context-dependent suppression of termination codons in a mammalian translation system. RNA 6, 1044–1055.

Mendell, J.T., Sharifi, N.A., Meyers, J.L., Martinez-Murillo, F., and Dietz, H.C. (2004). Nonsense surveillance regulates expression of diverse classes of mammalian transcripts and mutes genomic noise. Nat Genet 36, 1073–1078.

Mills, E.W., and Green, R. (2017). Ribosomopathies: There’s strength in numbers. Science 358.

Mills, E.W., Wangen, J., Green, R., and Ingolia, N.T. (2016). Dynamic Regulation of a Ribosome Rescue Pathway in Erythroid Cells and Platelets. Cell Rep 17, 1–10.

Mohn, F., Buhler, M., and Muhlemann, O. (2005). Nonsense-associated alternative splicing of T-cell receptor beta genes: no evidence for frame dependence. RNA 11, 147–156.

Muhlemann, O., and Lykke-Andersen, J. (2010). How and where are nonsense mRNAs degraded in mammalian cells? RNA Biol 7, 28–32.

Muhlrad, D., and Parker, R. (1999). Aberrant mRNAs with extended 3 ‘UTRs are substrates for rapid degradation by mRNA surveillance. RNA 5, 1299–1307.

Nagy, E., and Maquat, L.E. (1998). A rule for termination-codon position within intron-containing genes: when nonsense affects RNA abundance. Trends Biochem Sci 23, 198–199.

Neu-Yilik, G., Raimondeau, E., Eliseev, B., Yeramala, L., Amthor, B., Deniaud, A., Huard, K., Kerschgens, K., Hentze, M.W., Schaffitzel, C., et al. (2017). Dual function of UPF3B in early and late translation termination. EMBO J.

Nicholson, P., Gkratsou, A., Josi, C., Colombo, M., and Muhlemann, O. (2018). Dissecting the functions of SMG5, SMG7, and PNRC2 in nonsense-mediated mRNA decay of human cells. RNA 24, 557–573.

Peixeiro, I., Inacio, A., Barbosa, C., Silva, A.L., Liebhaber, S.A., and Romao, L. (2011). Interaction of PABPC1 with the translation initiation complex is critical to the NMD resistance of AUG-proximal nonsense mutations. Nucleic Acids Res 40, 1160–1173.

Pisarev, A.V., Skabkin, M.A., Pisareva, V.P., Skabkina, O.V., Rakotondrafara, A.M., Hentze, M.W., Hellen, C.U., and Pestova, T.V. (2010). The role of ABCE1 in eukaryotic posttermination ribosomal recycling. Mol Cell 37, 196–210.

Pulak, R., and Anderson, P. (1993). mRNA surveillance by the Caenorhabditis elegans smg genes. Genes Dev 7, 1885–1897.

Roy, B., Leszyk, J.D., Mangus, D.A., and Jacobson, A. (2015). Nonsense suppression by near-cognate tRNAs employs alternative base pairing at codon positions 1 and 3. Proc Natl Acad Sci U S A 112, 3038–3043.

Schmidt, E.K., Clavarino, G., Ceppi, M., and Pierre, P. (2009). SUnSET, a nonradioactive method to monitor protein synthesis. Nat Methods 6, 275–277.

Serdar, L.D., Whiteside, D.L., and Baker, K.E. (2016). ATP hydrolysis by UPF1 is required for efficient translation termination at premature stop codons. Nature communications 7, 14021.

Shoemaker, C.J., and Green, R. (2011). Kinetic analysis reveals the ordered coupling of translation termination and ribosome recycling in yeast. Proc Natl Acad Sci U S A 108, E1392–1398.

Shoemaker, C.J., and Green, R. (2012). Translation drives mRNA quality control. Nat Struct Mol Biol 19, 594–601.

Silva, A.L., Ribeiro, P., Inacio, A., Liebhaber, S.A., and Romao, L. (2008). Proximity of the poly(A)-binding protein to a premature termination codon inhibits mammalian nonsense-mediated mRNA decay. RNA 14, 563–576.

Singh, G., Rebbapragada, I., and Lykke-Andersen, J. (2008). A competition between stimulators and antagonists of Upf complex recruitment governs human nonsense-mediated mRNA decay. PLoS Biol 6, e111.

Stalder, L., and Muhlemann, O. (2008). The meaning of nonsense. Trends Cell Biol 18, 315–321.

Sudmant, P.H., Lee, H., Dominguez, D., Heiman, M., and Burge, C.B. (2018). Widespread Accumulation of Ribosome-Associated Isolated 3’ UTRs in Neuronal Cell Populations of the Aging Brain. Cell Rep 25, 2447–2456 e2444.

Tani, H., Imamachi, N., Salam, K.A., Mizutani, R., Ijiri, K., Irie, T., Yada, T., Suzuki, Y., and Akimitsu, N. (2012). Identification of hundreds of novel UPF1 target transcripts by direct determination of whole transcriptome stability. RNA Biol 9, 1370–1379.

Thermann, R., Neu-Yilik, G., Deters, A., Frede, U., Wehr, K., Hagemeier, C., Hentze, M.W., and Kulozik, A.E. (1998). Binary specification of nonsense codons by splicing and cytoplasmic translation. EMBO J 17, 3484–3494.

Toompuu, M., Karblane, K., Pata, P., Truve, E., and Sarmiento, C. (2016). ABCE1 is essential for S phase progression in human cells. Cell Cycle 15, 1234–1247.

Viegas, M.H., Gehring, N.H., Breit, S., Hentze, M.W., and Kulozik, A.E. (2007). The abundance of RNPS1, a protein component of the exon junction complex, can determine the variability in efficiency of the Nonsense Mediated Decay pathway. Nucleic Acids Res 35, 4542–4551.

Wittmann, J., Hol, E.M., and Jack, H.M. (2006). hUPF2 Silencing Identifies Physiologic Substrates of Mammalian Nonsense-Mediated mRNA Decay. Mol Cell Biol 26, 1272–1287.

Yan, X., Hoek, T.A., Vale, R.D., and Tanenbaum, M.E. (2016). Dynamics of Translation of Single mRNA Molecules In Vivo. Cell 165, 976–989.

Yepiskoposyan, H., Aeschimann, F., Nilsson, D., Okoniewski, M., and Muhlemann, O. (2011). Autoregulation of the nonsense-mediated mRNA decay pathway in human cells. RNA 17, 2108–2118.

Young, D.J., Guydosh, N.R., Zhang, F., Hinnebusch, A.G., and Green, R. (2015). Rli1/ABCE1 Recycles Terminating Ribosomes and Controls Translation Reinitiation in 3’UTRs In Vivo. Cell 162, 872–884.

Zuccotti, P., and Modelska, A. (2016). Studying the Translatome with Polysome Profiling. Methods Mol Biol 1358, 59–69.

